# Mucin *O*-glycans facilitate symbiosynthesis to maintain gut immune homeostasis

**DOI:** 10.1101/655597

**Authors:** Takahiro Yamada, Shingo Hino, Hideki Iijima, Tomomi Genda, Ryo Aoki, Ryuji Nagata, Ho Han, Masato Hirota, Yusuke Kinashi, Hiroyuki Oguchi, Wataru Suda, Yukihiro Furusawa, Yumiko Fujimura, Jun Kunisawa, Masahira Hattori, Michihiro Fukushima, Tatsuya Morita, Koji Hase

**Affiliations:** Division of Biochemistry, Faculty of Pharmacy, Keio University, Minato-ku, Tokyo, Japan; Department of Applied Biological Chemistry, Graduate School of Agriculture, Shizuoka University, Shizuoka, Japan; Department of Gastroenterology and Hepatology, Graduate School of Medicine, Osaka University, Osaka, Japan; Division of Gastroenterology and Hepatology, School of Medicine, Keio University, Shinjuku-ku, Tokyo, Japan; Department of Food Science, Obihiro University of Agriculture and Veterinary Medicine, Hokkaido, Japan; Graduate School of Frontier Sciences, The University of Tokyo, Chiba, Japan; Department of Microbiology and Immunology, Keio University School of Medicine, Tokyo, Japan; Department of Liberal Arts and Sciences, Toyama Prefectural University, Toyama, Japan; Laboratory of Vaccine Materials and Laboratory of Gut Environmental System, National Institutes of Biomedical Innovation, Health and Nutrition (NIBIOHN), Osaka, Japan; Department of Microbiology and Immunology, Kobe University Graduate School of Medicine, Hyogo, Japan; Graduate School of Medicine, Graduate School of Pharmaceutical Sciences, Graduate School of Dentistry, Osaka University, Osaka, Japan; International Research and Development Center for Mucosal Vaccines, The Institute of Medical Science, The University of Tokyo (IMSUT), Tokyo, Japan; Graduate School of Advanced Science and Engineering, Waseda University, Tokyo, Japan

## Abstract

The dysbiosis of gut microbiota has been implicated in the pathogenesis of inflammatory bowel diseases (IBDs); however, the underlying mechanisms have not yet been elucidated. Heavily glycosylated mucin not only establishes a first-line barrier against pathogens, but also serves as a niche for microbial growth. We hypothesized that dysbiosis may cause abnormal mucin utilization and microbial metabolic dysfunction. To test this hypothesis, we analyzed short-chain fatty acids (SCFAs) and mucin components in the stool samples of 40 healthy subjects, 49 ulcerative colitis (UC) patients, and 44 Crohn’s disease (CD) patients from Japan. The levels of *n*-butyrate were significantly lower in the stools of both the CD and UC patients than in those of the healthy subjects. Correlation analysis identified 7 bacterial species positively correlated with *n*-butyrate levels, among which the major *n*-butyrate producer, *Faecalibacterium prausnitzii*, was particularly underrepresented in CD patients, but not in UC patients. In UC patients, there were inverse correlations between mucin *O*-glycan levels and the production of SCFAs, such as *n*-butyrate, suggesting that mucin *O*-glycans act as an endogenous fermentation substrate for *n*-butyrate production. Indeed, mucin-fed rodents exhibited enhanced *n*-butyrate production, leading to the expansion of RORgt*^+^*Treg cells and IgA-producing cells in the colonic lamina propria. Importantly, the availability of mucin-associated *O*-glycans to the microbiota was significantly reduced in *n*-butyrate-deficient UC patients. Taken together, our findings highlight the biological significance of the symbiosynthesis pathway in the production of *n*-butyrate, which maintains gut immune homeostasis.

## Introduction

Inflammatory bowel diseases (IBDs), such as ulcerative colitis (UC) and Crohn’s disease (CD), are recurrent inflammatory disorders caused by both genetic and environmental factors [1,2]. Accumulating evidence has demonstrated that abnormal gut microbial composition, termed dysbiosis, plays a role in the pathogenesis and/or exacerbation of IBD in Caucasian patients [3,4]. The gut microbiota of IBD patients is characterized by diminished microbial diversity alongside the underrepresentation of Firmicutes and overrepresentation of Proteobacteria [3,5–8]. These characteristics are evident in the gut microbiota of CD patients, and the severity of dysbiosis in rectal mucosa-associated microbiota correlates well with disease score [7]. Changes in gut microbiota composition are less obvious in UC patients that in CD patients, and their association with IBD pathogenesis has not yet been elucidated [6,8]. Animal experiments have demonstrated that the gut microbiota shapes the host intestinal immune system under physiological conditions by inducing the maturation of gut-associated lymphoid tissues and the differentiation of Th17 and regulatory T cells [9]. In contrast, the gut microbiota drives intestinal inflammation under dysbiosis [10,11]. Similarly, the transplantation of microbiota from CD patients into *Il10*-deficient mice has been shown to accelerate colitis [12]; however, the underlying mechanisms by which dysbiosis promotes the inflammatory response have not yet been fully elucidated.

Short-chain fatty acids (SCFAs) such as acetate, propionate, and *n*-butyrate, are produced by intestinal microbiota via the microbial fermentation of indigestible carbohydrates [i.e. soluble dietary fibers, oligosaccharides, and resistant starches (RS)] in the colon and cecum of humans [13] and rodents [14], respectively. SCFAs not only serve as nutrients for the colonic epithelium [15], but also enhance mucosal barrier function by maintaining epithelial integrity [16], increasing mucin production [17], and triggering the IgA response [18]. Furthermore, *n*-butyrate exhibits anti-inflammatory effects by suppressing NF-κB signaling in the colonic epithelium [19]. It has previously been reported that *n*-butyrate facilitates regulatory T cell differentiation in the colonic lamina propria [20,21]. *n*-Butyrate is mainly produced by bacterial species classified under *Clostridium* cluster IV or XIVa. In IBD patients, these butyrate producers, including *Faecalibacterium prausnitzii* (*Clostridium* cluster IV) and Lachnospiraceae (*Clostridium* cluster XIVa) [22,23], are markedly underrepresented compared to that in healthy subjects [24–26]. SCFA mixture or *n*-butyrate enemas have been shown to effectively reduce the inflammatory symptoms of UC on the colonic mucosa [27,28]. These observations suggest that reduced *n*-butyrate production in IBD patients may exacerbate gut inflammation.

The luminal secretion of mucins by goblet cells establishes a mucus layer which functions as a physicochemical barrier, thus preventing microbial attachment to the epithelial surface [29]. Mucins predominantly consist of the heavily glycosylated MUC2 protein, which confers viscosity to the mucus in order to restrict bacterial motility. The stratified inner layer of the mucus is impenetrable to intestinal bacteria [30], whereas the outer layer of the mucus serves as a habitat for certain bacterial species which utilize mucin as an energy resource [31,32]. Experimental observations using mice devoid of Muc2 or its *O*-glycans have demonstrated that mucin barrier dysfunction enhances bacterial translocation into the lamina propria, resulting in chronic inflammation of the colon [33,34]. The same has been shown for UC, since the colonic mucus of UC patients is less protective than that of healthy subjects, therefore commensal bacteria often penetrate into the inner mucus layer [35].

The aforementioned abnormalities in mucin and SCFA production due to dysbiosis may be implicated in the development of IBD; however, the relevance of these major pathological events remains obscure. Therefore, we explored the relationship between the intestinal microbiota, SCFA levels, and mucin profiles of Japanese IBD patients. We observed reduced *n*-butyrate levels in the stools of both CD and UC patients; however, this may have been a result of different etiologies. Considerable reductions in the number of *n*-butyrate-producing bacteria diminished *n*-butyrate synthesis in CD patients, whereas reduced mucin *O*-glycan availability was observed in UC patients lacking *n*-butyrate. Herein, we demonstrated that mucin *O*-glycan is utilized by intestinal microbiota as an endogenous fermentation source to produce *n*-butyrate, and that this pathway is likely affected in UC.

## Results

### Reduced n-butyrate in the stools of IBD patients

To accurately characterize the altered composition of the intestinal microbiota in IBD patients, we initially analyzed the concentrations of various organic acids, including SCFAs, in stool samples. The disease status of UC and CD patients was classified as either active or in remission based on endoscopic assessment (S1 and S2 Table). We observed that fecal *n*-butyrate levels were significantly lower in the UC and CD groups than that in the healthy subjects (Fig 1). Notably, a substantial number of CD and UC patients lacked *n*-butyrate in their stools, particularly patients with active CD, 50 % of whom were devoid of *n*-butyrate. Thus, reduced stool *n*-butyrate levels reflect disease status in CD. In contrast, *n*-butyrate concentration was not affected by any other factors, including the site of disease development (i.e. right or left colon in UC, small or large intestine in CD), treatment with probiotics (lactic acid bacteria and *Clostridium butyricum*), 5-azathiopurine, and anti-TNF antibodies (S2 Fig, S3 and S4 Table).

**Fig 1.**
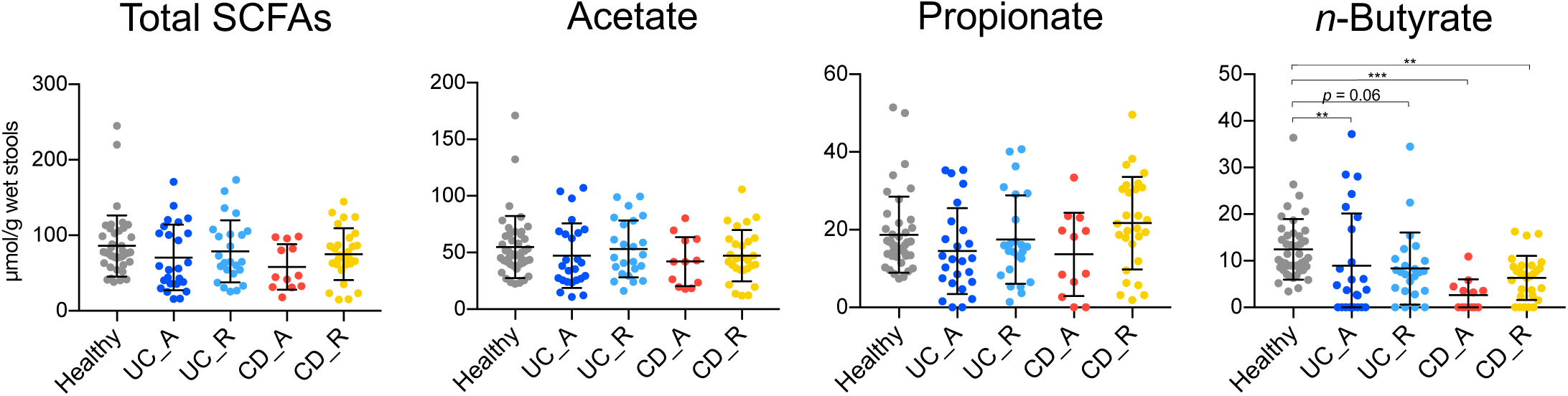
Fecal organic acid concentrations in IBD patients and healthy subjects. Fecal organic acid concentrations were analyzed in healthy subjects, patients with active (UC-A) or remissive (UC-R) UC, and patients with active (CD-A) or remissive (CD-R) CD. Data represent the mean ± SD (n = 12-43/group). \**p* < 0.05, \*\**p* < 0.01, and \*\*\**p* < 0.001 (ANOVA followed by Tukey’s multiple comparison test or the Kruskal-Wallis test followed by Dunn’s multiple comparison test).

The stool concentrations of propionate, another beneficial metabolite [36], were also slightly decreased in active UC and CD, although these differences were not statistically significant. *n*-Valerate was detected in 16.3 % of the healthy subjects, but not in UC patients (S1A Fig), whereas elevated levels of succinate were observed in a small number of UC and CD patients. The amounts of other organic acids, including acetate, formate, and lactate, were similar among the groups. Correlation analysis between the individual organic acids revealed that *n*-butyrate levels were positively correlated with propionate and acetate, but not formate, lactate, or succinate (S1B Fig). Together, these observations support the notion that reduced levels of anti-inflammatory SCFAs, especially *n*-butyrate, may be involved in the development and/or exacerbation of IBDs.

### Characterization of microbial communities in IBD patients

We performed 16S rRNA sequencing on the stool samples to characterize the microbial community composition of the IBD patients. The obtained data were subjected to principal coordinate analysis (PCoA) based on unweighted UniFrac (Fig 2A). The microbiota of the healthy subjects were grouped within one region, whereas those of the UC and CD patients were scattered, indicating that the microbial compositions of UC and CD patients were distinct from those of the healthy subjects (adonis based on unweighted UniFrac versus healthy microbiota; UC: *R*^2^ = 0.074 and *p* = 0.001; CD: *R*^2^ = 0.18 and *p* = 0.001). Furthermore, microbial composition differed significantly different between the active and remission phases of both diseases (S3 Fig). In accordance with previous studies of Caucasian IBD patients [6,7], α-diversity was significantly reduced in the Japanese UC and CD patients, particularly during the active stages (Fig 2B).

**Fig 2.**
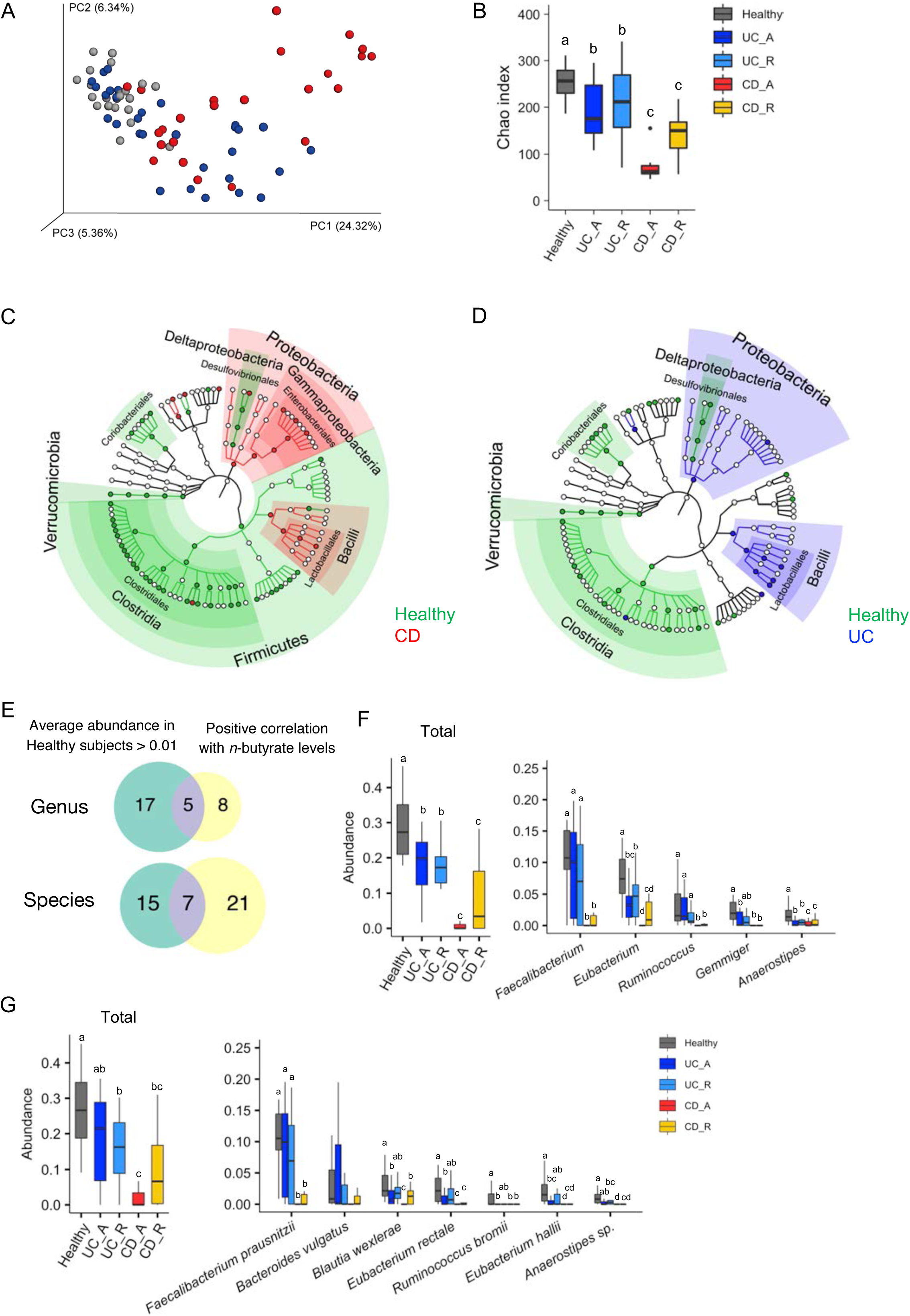
Microbial signature of IBD patients and healthy subjects. (A) Principal coordinate analysis of unweighted UniFrac distance among the microbiota of IBD patients and healthy subjects. (B) Species richness (Chao 1) of the microbiota of IBD patients and healthy subjects. (C, D) Taxa discriminating between the microbiota of (C) CD or (D) UC patients and healthy subjects, determined by LEfSe [38] analysis. The discriminating taxa for the indicated groups are annotated on the phylogenetic trees. (E) Venn diagram showing the criteria for butyrate-associated bacteria. (F, G) Relative abundance of butyrate-associated bacteria at the genus (F) and species (G) levels. The left panels represent the total number of butyrate-associated bacteria, and the right panels indicate the abundance of individual bacterial genera or species. Each boxplot represents the median, interquartile range (IQR), and the lowest and highest values within 1.5 IQRs of the first and third quartiles (n = 7-23/group). The outliers are not shown. Different letters over the boxplots indicate significant differences (*p* < 0.05; Kruskal-Wallis test followed by Dunn’s multiple test).

The microbiota of the CD patients was characterized by the underrepresentation of Firmicutes and Verrucomicrobia (including only *Akkermansia muciniphila*), and the overrepresentation of Proteobacteria at the phylum level (Fig 2C and S4A Fig). The latter was also slightly elevated in UC patients. At the class level, patients with active CD exhibited decreased occupancy of Clostridia and a reciprocal increase in the occupancy of Gammaproteobacteria and Bacilli (S4B Fig). BugBase [37] predicted the outgrowth of facultative anaerobes and the reduction of anaerobes in this group (S5 Fig). A similar trend was also observed to a lesser extent in the microbiota of CD patients in remission, active UC patients, and UC patients in remission (Fig 2A-D, S4 and S5 Fig).

### Reduced numbers of butyrate-associated bacteria in IBD patients

To identify the bacterial species contributing towards the production of intestinal metabolites, including *n*-butyrate, we performed correlation analyses between the abundance of bacterial taxa and each metabolite. We found that the abundance of 12 genera and 28 species were positively correlated with *n*-butyrate levels (Spearman correlation analysis, FDR < 0.05; S5 and S6 Table), whereas only a few genera were correlated with acetate, *n*-valerate, or *i*-butyrate levels. The other metabolites analyzed here showed no significant correlation with any specific genera or species. Among the *n*-butyrate-associated bacteria, 5 bacterial genera and 7 species occupied 1 % or more of the microbial communities of healthy subjects, suggesting that these bacteria may play a central role in *n*-butyrate biosynthetic pathways (Fig 2E). Thereafter, we termed these “butyrate-associated bacteria” (Fig 2F and 2G), which included well-known *n*-butyrate producers such as *F. prausnitzii* and *Eubacterium rectale* [22]. Consistent with the data concerning stool *n*-butyrate concentration, the overall abundance of butyrate-associated bacteria was considerably lower in CD patients (Fig 2F and 2G). Notably, *F. prausnitzii*, which was the most abundant *n*-butyrate producer in the microbiota of healthy individuals, was almost absent in the microbiota of active CD patients (Fig 2G, right). In contrast, the abundance of *F. prausnitzii* was not significantly reduced in active or remission UC patients (Fig 2F and 2G). The minor populations of *n*-butyrate-associated bacteria, namely, *Blautia wexlerae, Eubacterium rectale, Ruminococcus bromii*, and *E. hallii* were underrepresented in active UC patients (Fig 2G, right).

### Reduced availability of mucin O-glycans to UC-associated microbiota

Mucin *O*-glycans form complex structures, consisting of *N*-acetylgalactosamine, galactose, and *N*-acetylglucosamine, which are elongated and terminated by fucose, sialic acids, and sulfate residues in the human colon [39]. We hypothesized that the quality and quantity of the mucin components may be affected in IBD. We therefore explored the amounts of these components in the stool mucin (Fig. 3). The protein content of the mucin fraction was significantly lower in the UC patients, which may reflect may reflect the previously reported downregulation of Muc2 expression [40,41]. Nevertheless, mucin *O*-glycan levels were slightly higher in these patients than in the healthy subjects (Fig. 3B). Consequently, the mucin ratio of *O*-glycans to protein was significantly higher in the UC patients, indicating the reduced availability of mucin *O*-glycans to UC-associated microbiota. This abnormality was characteristic of UC, but not of CD. Notably, the composition of mucin *O*-glycan components was mostly intact in UC and CD patients (S7 Table), except for NeuGc which was detected at higher levels in the UC patients than in the healthy subjects (S8 Table).

**Fig 3.**
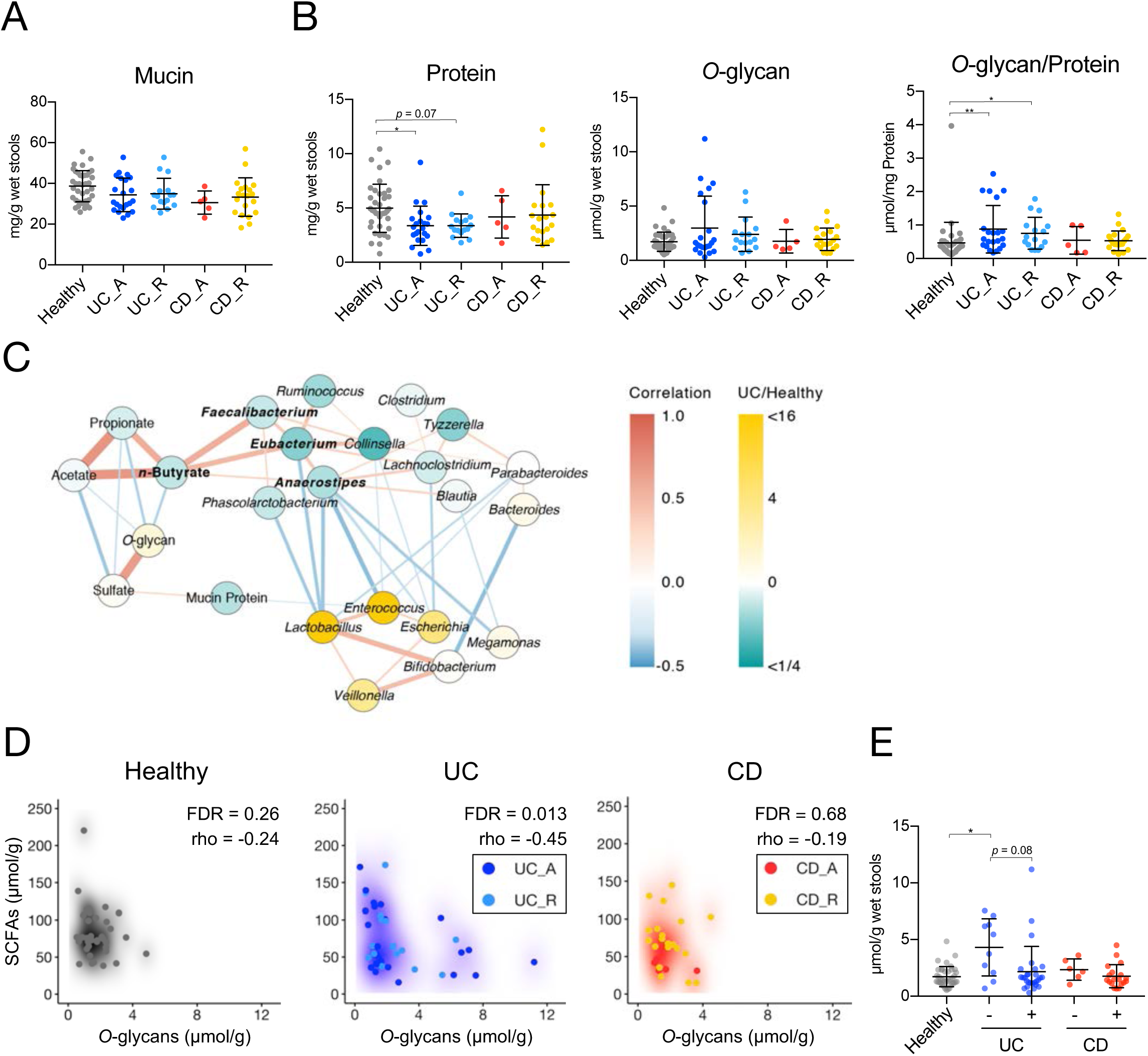
Stool mucin components of IBD patients and healthy subjects. (A, B) The levels of mucin (A), mucin-associated protein, and (B), *O*-glycan were analyzed in the stools of the IBD patients and healthy subjects. Data represent the mean ± SD (n = 5-38/group). \**p* < 0.05 and \*\**p* < 0.01 (ANOVA followed by Tukey’s multiple comparison test or the Kruskal-Wallis test followed by Dunn’s multiple comparison test). (C) Correlation network of the levels of SCFAs, mucin components, and bacteria in the stool samples. Nodes represent the SCFAs, mucin components, or bacterial genera with average levels higher than 1.0 %. Node colors reflect the average levels in UC patients compared to those in healthy subjects. Red and blue lines indicate positive (Spearman’s correlation coefficient > 0.3) and negative (correlation coefficient < –0.3) correlations, respectively. Edge thickness represents the strength of the correlation. Line color intensity reflects the extent of the correlation. (D) Scatter density plots of fecal *n*-butyrate levels versus mucin *O*-glycan levels for individual disease groups. The increase in the intensity of each color (white to black, blue, or red) reflects the density of the scatter plot. Spearman coefficients (rho) and FDRs are shown. (E) UC and CD patients were classified into two groups each based on fecal *n*-butyrate detection in order to compare fecal levels of mucin *O*-glycans. Patients with *n*-butyrate concentrations below 0.1 μmol/g were classified as “Undetected”, while the others were classified as “Detected”.

### Impaired mucin O-glycan availability may reduce n-butyrate production in UC patients

We generated correlation networks (rho > 0.3 or rho < –0.3) of bacterial genera, mucin components, and SCFAs. Correlation network analysis highlighted positive correlations between *n*-butyrate concentration and butyrate-associated bacteria (i.e. *Faecalibacterium*, *Eubacterium*, and *Anaerostipes*) (Fig 3C). The butyrate-associated bacteria were inversely correlated with *Escherichia*, *Enterococcus*, and *Lactobacillus*, which were overrepresented in UC patients (S6 Fig). Notably, network analysis revealed inverse correlations between the levels of mucin *O*-glycan and each SCFA (Fig 3C). Of the three groups, the inverse correlations between mucin *O*-glycan and the SCFAs were the most prominent in UC patients (total SCFAs: rho = –0.45, FDR = 0.014; acetate: rho = –0.44, FDR = 0.014; propionate: rho = –0.43, FDR = 0.014; *n*-butyrate: rho = –0.35, FDR = 0.035; Fig 3D and S5 Fig). These data suggest that intestinal microbiota could utilize mucin *O*-glycans as an endogenous fermentation substrate to produce SCFAs, such as *n*-butyrate. This idea was further supported by the reduced utilization of mucin *O*-glycans in the *n*-butyrate-deficient UC patients (Fig 3D). Thus, the development of dysbiosis may impair mucin *O*-glycan availability in patients with UC, which could eventually affect *n*-butyrate production.

To confirm this hypothesis, we assessed mucin glycan degradation (mucinase activity) in the fresh stool samples of newly recruited UC patients and healthy subjects (Fig 4). We observed a clear decrease in the stool mucinase activity in the UC patients compared to that in the healthy subjects. Mucinase activity exhibited a positive correlation with the stool concentrations of total SCFAs and butyrate, both of which were significantly lower in the UC patients (Fig 4B and S9 Fig). The decrease in SCFA levels was not attributed to the dietary habits of the UC patients since there were no significant differences in the nutrient intake (including soluble fiber intake) of the UC patients and healthy subjects (S10 Fig and S9 Table). Taken together, these data suggest that abnormal mucin *O*-glycan utilization may cause reduced *n*-butyrate production in UC.

**Fig 4.**
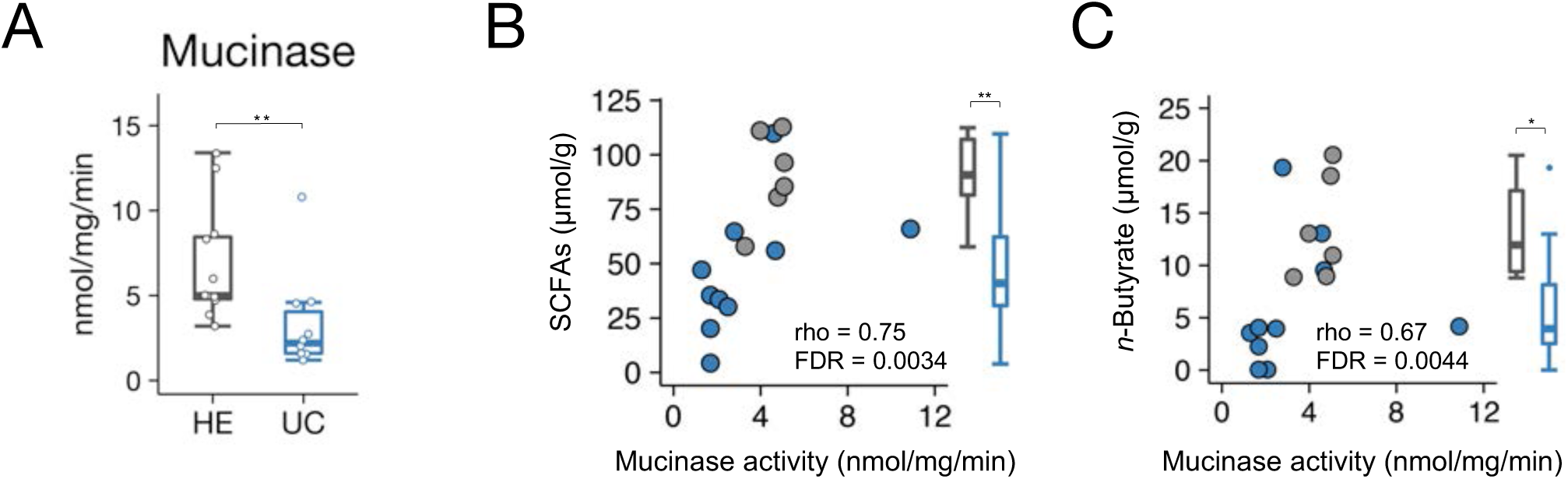
Stool mucinase activity in UC patients and healthy subjects. (A) The stool samples of UC patients and healthy subjects were analyzed for mucinase activity. Boxplots represent the median, interquartile range (IQR), and the lowest and highest values within 1.5 IQRs of the first and third quartiles (*n* = 10-11/group). (B, C) Scatter plots of mucinase activity and (B) total SCFAs and (C) *n*-butyrate concentrations. Spearman’s correlation coefficient (rho) and FDR are shown. Box plots represent stool concentrations of total SCFAs (B) and butyrate (C) in the two groups (*n* = 6-10/group). \**p* < 0.05 and \*\**p* < 0.01 (Wilcoxon’s rank sum test).

### Mucin supplementation facilitates SCFA production

To directly investigate whether the gut microbiota utilize mucin as a fermentation source to produce SCFAs, we cultured rat cecal microbiota in the presence or absence of mucin using an *in vitro* fermentation system [42]. The cecum is a major site of microbial fermentation in rodents [43]. Within 48 h, mucin supplementation had increased the concentration of SCFAs, including *n*-butyrate, in the culture media (Fig 5A). Thus, the microbial fermentation of mucin *O*-glycans generates SCFAs.

**Fig 5.**
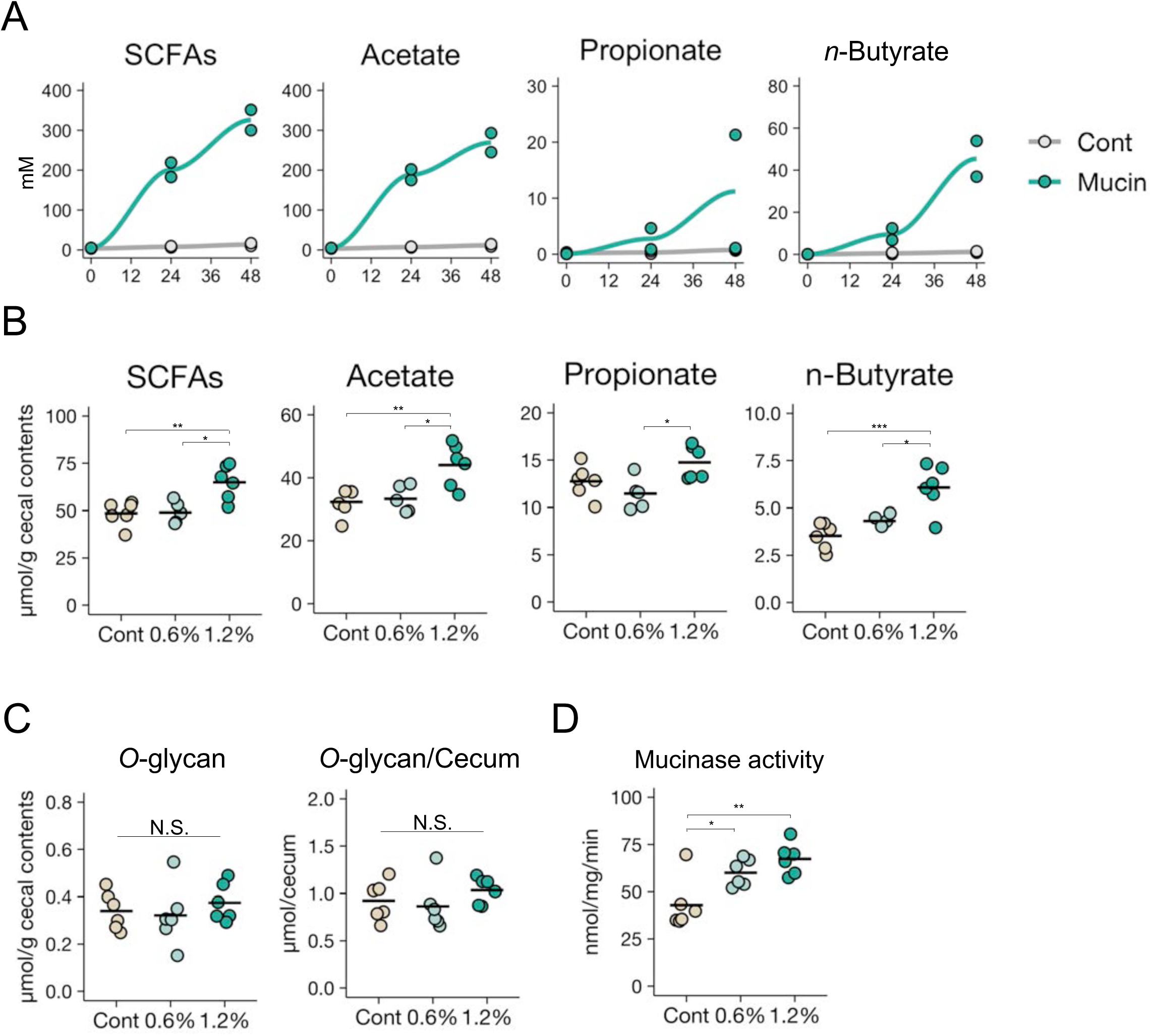
Mucin supplementation increases SCFA production by gut microbiota. (A) Rat cecal contents were cultured in the *in vitro* fermentation system in the presence or absence of porcine gastric mucin and SCFA concentrations were measured at the indicated time points. Data are representative of two independent experiments with similar results. (B-D) Cecal SCFA levels (B), *O*-glycan levels (C), and mucinase activity (D) in the stools of rats fed a control diet or diets containing 0.6 or 1.2 % porcine stomach mucin. Data represent the mean ± SD (*n* = 5-6). \**p* < 0.05 and \*\**p* < 0.01 (ANOVA followed by Dunnett’s multiple comparison test or the Kruskal-Wallis test followed by Dunn’s multiple comparison test, compared to the control group).

We further corroborated the importance of mucin *O*-glycans as an endogenous fermentation substrate *in vivo*. We fed rats synthetic diets containing 0.6 or 1.2 % (w/w) purified mucins for 2 weeks. The 1.2 % mucin diet significantly increased acetate and *n*-butyrate concentrations in the ceca (Fig 5B), and slightly increased the propionate concentration. Importantly, the level of mucin *O*-glycans in the cecum was similar in all the groups (Fig 5C), indicating that the exogenous mucin *O*-glycans were efficiently consumed via intestinal microbial fermentation. Furthermore, mucin administration increased microbial mucinase activity in a dose-dependent manner (Fig 5D). Thus, *O*-glycan utilization may be a key determinant for maintaining SCFA production.

### The mucin-SCFA axis shapes the gut immune system

Finally, we investigated the biological significance of the mucin-dependent symbiosynthetic pathway by feeding mice a 1.5 % mucin-containing diet for 3 weeks. The microbiota of the mucin-fed mice was considerably different to that of the controls, characterized by higher α-diversity and increased abundances of *Allobaculum*, unclassified Bacteroidales S24-7, and *Akkermansia* (S11 Fig). Consistent with the observations made in the rats, the administration of mucin to mice increased the cecal concentration of SCFAs such as *n*-butyrate (Fig 6A). Gut microbiota-derived butyrate induces the differentiation of peripherally generated Treg (pTreg) cells [18,20,44], while SCFAs facilitate the development of B220^−^IgA^+^ plasma cells in the colonic lamina propria [18,20,44]. In agreement with these reports, the frequencies of RORγt^+^Foxp3^+^ pTreg cells in the mucin-fed mice was twice as high as in the control mice (Fig 6B and 6C). Notably, the mucin diet also increased RORγt^−^Foxp3^+^ thymus-derived Treg (tTreg) cells, suggesting that the colonic migration and/or proliferation of tTreg cells was also enhanced in the mucin-fed group. Furthermore, the mucin-containing diet increased the number of IgA^+^ plasma cells in the colon (Fig 6D and 6E). Based on these observations, mucin not only serves as a mucosal barrier but also shapes the intestinal immune system by facilitating microbial fermentation to increase luminal SCFA concentrations (S12 Fig.).

**Fig 6.**
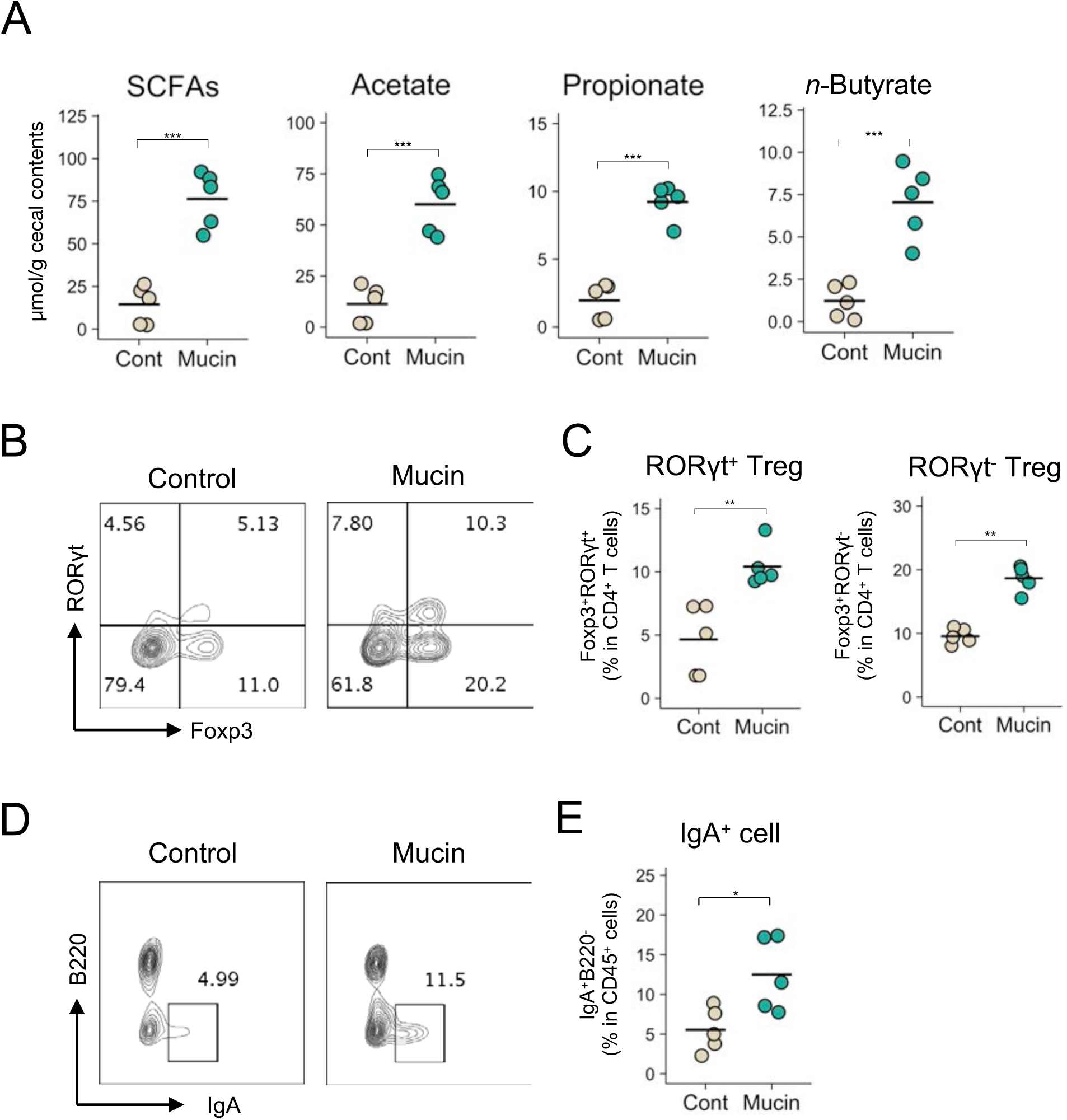
Immune modulation by the mucin-containing diet. (A-E) Cecal SCFA levels (A), the frequency of Foxp3^+^RORγt^+^ cells and Foxp3^+^RORγt^−^ cells within CD4^+^ T cells (B and C), and IgA^+^B220^−^ cells within CD45^+^ cells (D and E) in the colonic mucosa of mice fed a control diet or a diet containing 1.5 % porcine stomach mucin for 3 weeks. Representative flow cytometry plots are shown in B and D. Data represent the mean ± SD (*n* = 5). \**p* < 0.05, \*\**p* < 0.01, and \*\*\**p* < 0.001 (two-tailed Student’s *t*-test).

## Discussion

In this study, we confirmed that Japanese IBD patients exhibit symptoms of dysbiosis, and that the intestinal microenvironments of CD and UC patients display several different features. The microbial community was less diverse in CD patients than in UC patients (Fig 2B), whilst microbial composition during the active stage of CD was characterized by the expansion of Gammaproteobacteria and Bacilli alongside an underrepresentation of Clostridia (Fig 2C). Consequently, the frequency of anaerobes was markedly decreased (S5 Fig), and *n*-butyrate producers (mainly *F. prausnitzii*) were nearly absent during active CD (Fig 2G). Diminished microbial diversity alongside an underrepresentation of *n*-butyrate producers has also been reported in Caucasian CD patients [7,8,24,26]; therefore, these microbial alterations are considered hallmarks of CD and are closely associated with disease severity. *N*-butyrate producers such as *F. prausnitzii* and *E. rectale* have been identified based on their *in vitro* metabolic activity [7,24–26]; however, the contribution of these bacteria to *n*-butyrate production in the human intestine has remained unclear. Our integrated analysis demonstrated that these bacteria are positively correlated with fecal *n*-butyrate concentration (Fig 2G), thus an underrepresentation of *n*-butyrate producers should be the main cause of reduced *n*-butyrate production in active CD. During CD remission, the abundances of *n*-butyrate-associated bacteria slightly increased compared to active CD, thus raising the fecal concentration of *n*-butyrate.

Conversely, the gut microbiota of UC patients showed only a slight decrease in microbial diversity and retained normal levels of *F. prausnitzii*, even during the active stage (Fig 2B and 2G). Nevertheless, some of the UC patients were deficient in fecal *n*-butyrate. To elucidate the underlying mechanism, we analyzed the mucin components which serve as a nutrient source for gut microbiota. The levels of mucin protein were lower in the stools of UC patients, presumably reflecting reduced net mucin production [40,41]. In contrast, the levels of mucin *O*-glycan in the stools were significantly higher in UC patients lacking stool *n*-butyrate than in healthy subjects. These observations imply that mucin *O*-glycan availability is compromised in the intestinal microbiota of UC patients. Indeed, the stool samples of UC patients exhibited lower mucinase activity than those of healthy subjects. Mucin *O*-glycan availability is considered a key determinant for the colony-forming ability of certain bacterial species on the mucosal surface [45,46]. Additionally, we believe that mucin *O*-glycans are utilized as an endogenous fermentation source to produce SCFAs. Several lines of evidence support this hypothesis. Firstly, there was an inverse correlation between the levels of mucin *O*-glycan and SCFAs in the stool samples of UC patients, whilst mucinase activity was positively correlated with the concentration of SCFAs. Secondly, it has previously been reported that substantial levels of SCFAs are still generated in the large intestines of rodents fed synthetic diets lacking dietary fiber and RS [43,47], indicative of the presence of endogenous fermentation sources in the gut. Intestinal microbiota may utilize mucin as a major fermentation source in the absence of fermentable dietary ingredients. We verified this hypothesis by showing that the exogenous administration of purified mucin facilitated the microbial production of SCFAs *in vitro* and *in vivo*. The mucin-containing diet increased the numbers of *Allobaculum*, unclassified Bacteroidales S24-7, and *Akkermansia* in the cecal microbiota of mice, suggesting that these bacteria may utilize mucin as an energy source to different degrees. Indeed, *Akkermansia* and certain *Bacteroides* spps. have been reported to degrade mucin [48,49]. Furthermore, mucinase activity was increased in the cecal contents of mucin-fed rats. *Allobacullum* species are known to produce butyrate; however, *Allobacullum* mainly consumes mono- or di-saccharides [50]. These results suggest that *Allobacullum* may utilize the mono- or di-saccharides produced from mucin *O*-glycans by the other two bacteria. Notably, the abundance of Lachnospiraceae and Ruminococcaceae, the major butyrate producers [22,23], was unchanged in the mucin-fed mice. Therefore, the administration of exogenous mucin is likely to have altered microbial composition by promoting the outgrowth of mucin utilizers and enhancing the production of SCFAs, including butyrate. Thirdly, previous studies using an *in vitro* gut model demonstrated that Firmicutes dominate the gut mucus microbiota, with *Clostridium* cluster XIVa species accounting for approximately 60 % of all mucus-colonizing bacteria [51]. 16S rRNA gene analysis of the mucosa-associated bacteria showed that Firmicutes were more abundant in the mucus region, with *Clostridium* cluster XIVa (Lachnospiraceae and Ruminococcaceae) being the predominant families [52]. Certain gut bacteria, such as *A. muciniphila*, are capable of utilizing mucin *O*-glycans, but not dietary polysaccharides [49]. Mucus-associated bacteria express mucin glycan utilization-associated genes at high levels, indicating that these bacteria utilize mucin glycans [32]. Therefore, we suspect that certain bacteria in the mucus utilize mucin *O*-glycans for *n*-butyrate production even in the presence of dietary fibers. Consequently, we propose a new model of symbiotic interaction in which host cell-derived mucin *O*-glycans are efficiently metabolized by intestinal microbiota under physiological conditions to produce *n*-butyrate, which is further utilized as an energy source by colonic epithelial cells and shapes the colonic immune system. This “symbiosynthesis” based on the mucin *O*-glycan–*n*-butyrate axis may play a significant role in establishing host–microbe symbiosis. In UC, the symbiosynthetic pathway seems to be impaired due to dysbiosis, at least partly contributing to the reduced production of *n*-butyrate which may be implicated in the pathogenesis of UC.

Furthermore, we demonstrated the immunomodulatory functions of mucin in the colon (Fig. 6B-E). We confirmed that exogenous mucin expanded Treg cell and IgA^+^ plasma cell populations, indicating that it has a significant role in shaping the immune system. Given that *n*-butyrate induces the differentiation of colonic pTreg cells and IgA^+^ plasma cells [18,20], we speculate that the intake of exogenous mucin increased these immune cell populations by enhancing *n*-butyrate biosynthesis. Interestingly, mucin administration also induced the expansion of tTreg cell populations (Fig. 6B and C), suggesting that mucin facilitates the differentiation, migration, and/or proliferation of tTreg cells. Increased propionate levels may be responsible for this event [53,54]. Although these mechanisms are not yet fully elucidated, these findings emphasize the importance of mucin as a fermentation source in the maintenance of colonic immune homeostasis.

Correlation analysis identified 7 species as butyrate-associated bacteria. Among these was the well-documented *n*-butyrate producer: *F. prausnitzii* [22]. *E. rectale* and *E. hallii*, which belong to the *Clostridium* cluster XIVa, also produce *n*-butyrate [22,23]. Interestingly, these *Eubacterium* spp. accumulate in the mucus layer [51,55] and their numbers are significantly lower in patients with active UC (Fig 2G). This may reflect the impaired mucin *O*-glycan–*n*-butyrate axis in UC. The genus *Anaerostipes* is another member of the *Clostridium* cluster XIVa and includes several butyrate producers, such as *A. caccae* and *A. hadrus* [56,57]. The remaining species, *R. bromii*, *B. wexlerae*, and *B. vulgatus*, may indirectly contribute to *n*-butyrate production as part of a bacterial metabolic web for *n*-butyrate biosynthesis. For example, *B. vulgatus* has an abundance of genes associated with polysaccharide utilization [58], whilst *R. bromii* and *B. wexlerae*, classified as *Clostridium* cluster IV and XIVa, respectively, are known as amylolytic bacteria. In particular, *R. bromii* plays a central role in the degradation and utilization of RS in the human colon [59] and *B. wexlerae* utilizes RS during fermentation *in vitro* [60]. Such carbohydrate digestion processes may be involved in *n*-butyrate production. *R. bromii* and *B. wexlerae* numbers were significantly lower in active UC patients; therefore, we speculate that the digestion of polysaccharides and RS might be reduced in the microbiota of these patients.

In conclusion, we observed that *n*-butyrate biosynthesis was decreased in both CD and UC patients; however, the underlying mechanisms differed between the two diseases (S12 Fig). In CD patients, *n*-butyrate levels are reduced by the loss of the majority of *n*-butyrate producers, including *F. prausnitzii*. Conversely, UC-associated microbiota utilize mucin-*O*-glycans poorly, which eventually leads to a decrease in *n*-butyrate production. Although further studies are required to clarify the precise molecular mechanisms of the mucin-*O*-glycan-dependent *n*-butyrate pathway, our findings offer a new perspective on the pathogenesis of UC and the development of intestinal dysbiosis (S12 Fig).

## Materials and Methods

### Sample collection

To analyze organic acid concentration, bacterial composition, and mucin components, the stool samples of 44 healthy subjects, 40 CD patients, and 49 UC patients from Japan were collected at the Department of Gastroenterology and Hepatology in Osaka University Hospital (Study 1; Table 1). In a separate study, the stool samples of 11 healthy subjects and 10 UC patients were collected to analyze mucinase activity (Study 2; Table 2). A validated food frequency questionnaire was used to investigate nutrient intake [61]. Patients were diagnosed with CD or UC according to endoscopic, radiological, histological, and clinical criteria provided by the Council for International Organizations of Medical Sciences of the World Health Organization and the International Organization for the Study of Inflammatory Bowel Disease [62–64]. Clinical disease activity was assessed via clinical disease activity and CDAI in UC and CD, respectively [65,66]. The endoscopic disease activity obtained by ileocolonoscopy was assessed using Matt’s score and modified Rutgeert’s score in UC and CD, respectively [67,68]. Endoscopic remission was defined as a Matt’s score of 1 in UC and a modified Rutgeert’s score of 0 or 1 in CD. All participants provided written informed consent to participate after receiving verbal and written information about the study. The protocol was approved by the ethics committees of Osaka University (#13165-2), The University of Tokyo (#25-42-1122), Keio University (#150421-1), Shizuoka University (#14-11), and NIBIOHN (#72) prior to subject inclusion.

**Table 1.** IBD patient information (Study 1). The information of the patient number, age, sex, disease duration, disease severity based on endoscopic and clinical diagnosis.

|  | Healthy | Ulcerative Colitis | Crohn's Disease |
| --- | --- | --- | --- |
|  | *n* = 44 | *n* = 49 | *n* = 40 |
| Median age, years (S.D.) | 33 (9.1) | 46 (13.8) | 40 (13.6) |
| Male (%) | 41 (95.3 %) | 27 (55.1 %) | 29 (72.5 %) |
| Disease Duration, years (S.D.) | - | 12.4 (10.5) | 13.4 (11.7) |
| Endoscopic Diagnosis |  |  |  |
| Active stage | - | 25 (51.0 %) | 12 (30 %) |
| Remission stage | - | 24 (49.0 %) | 28 (70 %) |
| Clinical Diagnosis |  |  |  |
| Active stage | - | 7 (14.3 %) | 10 (25 %) |
| Remission stage | - | 42 (85.7 %) | 30 (75 %) |

**Table 2.** UC patient information (Study 2). The information of the patient number, age, sex, disease duration, disease severity based on endoscopic and clinical diagnosis.

|  | Healthy | Ulcerative Colitis |
| --- | --- | --- |
|  | *n* = 11 | *n* = 10 |
| Mean age, years (S.D.) | 32.0 (12.7) | 42.8 (14.9) |
| Male (%) | 11 (100 %) | 6 (60 %) |
| Disease Duration, years (S.D.) | - | 10.3 (7.5) |
| Endoscopic Diagnosis |  |  |
| Active stage | - | 7 (70 %) |
| Remission stage | - | 3 (30 %) |
| Clinical Diagnosis |  |  |
| Active stage | - | 2 (20 %) |
| Remission stage | - | 8 (80 %) |

### Metagenomic 16S rRNA sequencing

Approximately 200 mg of the stool samples was transferred into 2 mL tubes containing 0.1 mm zirconia/silica beads (BioSpec Products, Bartlesville, OK) and 3.0 mm zirconia beads (Biomedical Sciences, Tokyo, Japan). The stool samples were homogenized at 1,500 rpm for 20 min with Shake Master Neo (Biomedical Sciences, Tokyo, Japan) after adding the Inhibit EX buffer from the QIAamp Fast DNA Stool Mini Kit (Qiagen, Hilden, Germany). Genomic DNA was then extracted using the kit according to the manufacturer’s instructions and was re-suspended in 10 mM Tris-HCl buffer at 5 ng/μL. A library of 16S rRNA genes was prepared according to the protocol described in an Illumina technical note [69]. Briefly, each DNA sample was amplified by polymerase chain reaction (PCR) using KAPA HiFi HS ReadyMix (KAPA Biosystems, Wilmington, MA) and primers specific for variable regions 3 and 4 of the 16S rRNA gene. The PCR products were purified using Agencourt AMPure XP Beads (Beckman Coulter, Brea, CA) and appended by PCR using the Nextera XT index kit (Illumina, San Diego, CA). The libraries were further purified using Agencourt AMPure XP Beads, diluted to 4 nM with 10 mM Tris-HCl buffer, and pooled. The pooled samples were sequenced using the Miseq system (Illumina) with a 2 × 300-base pair protocol. All the sequences analyzed in this study were deposited in the DNA Data Bank of Japan (DDBJ)/GenBank/European Molecular Biology Laboratory (EMBL) database under the accession number DRA006094.

### Bacterial composition analysis

We used the join_paired_ends.py QIIME script [70] and the fastq-join method to join paired-end reads, and trimmed sequencing adaptor sequences using cutadapt [71]. We converted the FASTQ files into FASTA files and removed chimera reads using the identify_chimeric_seqs.py and filter_fasta.py QIIME scripts with usearch61 [72]. Next, we concatenated the FASTA files of individual samples into one FASTA file and used the pick_open_reference_otus.py QIIME script to pick OTUs. We assigned taxonomy using the assign_taxonomy.py QIIME script with the RDP classifier and the Greengenes reference database, clustered at 97 % identity. Subsequently, we created an OTU table using the make_otu_table.py QIIME script and removed OTUs lower than 0.005 % using the filter_otus_from_otu_table.py script. We subsampled to a depth of 10,000 reads per sample and performed diversity analysis using the core_diversity_analyses.py QIIME script. To analyze the human stool microbiota, taxonomy was assigned to each OTU by similarity searching against the publicly available 16S (RDP ver. 10.27 and CORE update 2 September 2012) and NCBI genome databases using the GLSEARCH program, as described previously [73]. Phylum-, class-, genus-, and species-level assignments were performed at 70 %, 90 %, 94 %, and 97 % sequence identity thresholds, respectively. To analyze the cecal microbiota of the mice, taxonomic assignments were performed using the summarize_taxa.py QIIME script with the Greengenes reference database, clustered at 97 %, since there were abundant OTUs which lacked sequence homology with any species in the NCBI database. Diversity analyses of the fecal microbiota (α- and β-diversity) were performed using the Vegan package in R. The phenotype of the microbiome was predicted using BugBase [37] with the default parameters.

### Organic acid analysis

The levels of organic acids (formate, acetate, propionate, *i*-butyrate, *n*-butyrate, *i*-valerate, *n*-valerate, succinate, and lactate) in the human stools and rat cecal contents were measured according to the internal standard method using an HPLC (LC-6A; Shimadzu, Kyoto, Japan) equipped with a Shim-pack SCR-102H column (8 mm internal diameter × 30 cm long; Shimadzu) and an electroconductivity detector (CDD-6A; Shimadzu), as described previously [43]. Approximately 200 mg of human stool sample or 300 mg of rat cecal contents were homogenized in 2 mL of 10 mmol/L sodium hydroxide solution containing 0.5 g/L crotonic acid as an internal standard, then centrifuged at 10,000 x *g* for 15 min. The supernatant obtained was subjected to HPLC analysis. The detection limit for the organic acids was 0.1 μmol/g.

Levels of organic acids in the mouse cecal contents were measured using a gas chromatography-mass spectrometer (GC-MS) according to the modified methods of Moreau *et al.* [74]. Approximately 50 mg of mouse cecal contents was homogenized in 9 times the volume of H_2_O (w/w). After centrifugation (10,000 x g at 4 °C for 15 min), 200 μL of the supernatant was spiked with 10 µL of 1 mM 2-ethyl butyric acid (2-EB) as an internal standard and 20 µL of 20 % (w/v) 5-sulfosalichylic acid solution for deproteinization. After centrifugation (10,000 x g at 4 °C for 15 min), 200 μL of the supernatant was acidified using 10 μL of 37 % HCl, and organic acids were extracted via two rounds of 1 mL diethyl ether extraction. Next, 500 μL of the organic supernatant was mixed with 50 μL of *N*-tert-butyldimethylsilyl-*N*-methyltrifluoracetamide (MTBSTFA; Sigma-Aldrich Co., St. Louis, MO) in a new glass vial and left for 24 h at room temperature to derivatize. The derivatized samples were run through a JMS-Q1500GC GC/MS System (JEOL Ltd., Tokyo, Japan) equipped with an HP-5 capillary column (60 m × 0.25 mm × 0.25 μm, Agilent Technologies, Inc., Santa Clara, CA). Pure helium (99.9999 %) was used as a carrier gas and delivered at a flow rate of 1.2 mL/min. The following temperature program was used: 60 °C (3 min), 60–120 °C (5 °C/min), 120–290 °C (20 °C/min), 290 °C (3 min). Organic acid concentrations were determined by comparing their peak areas with the standards. Acetate was obtained from Nakarai Tesque, Inc., (Kyoto, Japan); propionate, *n*-butyrate, succinate and lactate from Wako Pure Chemical Industries, Ltd., (Osaka, Japan); and *i*-butyrate, *i*-valerate and *n*-valerate from Kanto Chemical Co., Inc (Tokyo, Japan).

### Mucin component analysis

The mucin fraction was isolated as described previously [75]. After being diluted appropriately, the *O*-linked oligosaccharide chains were analyzed as described previously [76]. Standard solutions of *N*-acetylgalactosamine (Sigma-Aldrich) were used to calculate the amount of oligosaccharide chains released from the mucins during the procedure. Approximately 20 mg of the mucin fraction was completely dried, resuspended in 200 mL of 4 mol/L HCl, and then hydrolyzed at 100 °C for 4 h in a heating block. The sulfate levels in the mucin fraction were determined according to the method of Harrison and Packer [77]. Solutions of 0.79, 1.59, 3.18, 6.38, and 12.8 mmol sulfate/L (Multi-anion standard solution-1; Wako Pure Chemicals, Osaka, Japan) were used as standards. Sialic acid levels were measured as described previously [78]. Briefly, the mucin fractions were hydrolyzed in 50 mM HCl at 80 °C for 1 h, and the *N*-acetylneuraminic acid (NeuAc) and *N*-glycolylneuraminic acid (NeuGc) released were labeled with 1,2-diamino-4,5-methylenedioxybenzene (DMB) using a commercial kit (Takara, Shiga, Japan). The DMB-labeled sialic acids were analyzed by HPLC equipped with a TSK-ODS80Ts column (Tosoh, Tokyo, Japan) and a fluorescence detector (RF-10AXL; Shimadzu).

### Preparation of mucin-containing diets

Semi-purified porcine stomach mucin (Sigma-Aldrich) was suspended in phosphate-buffered saline and filtered to remove contaminants. Ethanol was added to the filtrate to prepare a 60 % ethanol (w/w) solution. Glycosylated mucin was precipitated at −30 °C and collected by centrifugation. This treatment was repeated twice to further purify the mucin. The purified mucin was added to the AIN73-formula diet as a substitute for corn starch.

### Animal experiments

All animal experiments were performed according to protocols approved by the Animal Use Committees of Keio University and Shizuoka University. The rats and mice were maintained in accordance with the guidelines for the care of laboratory animals of Shizuoka University and Keio University, respectively. Male Wister rats were purchased from the Shizuoka Laboratory Animal Center and housed in individual wire screen-bottomed, stainless steel cages at 23 ± 2 °C in a lighting-controlled room (lit from 8:00 a.m. to 8:00 p.m.).

C57BL/6J male mice were purchased from CLEA Japan, Inc (Tokyo, Japan). The mice were fed an AIN93 diet (Oriental Yeast, Tokyo Japan) for 7 days, then either a 1.5 % mucin-containing diet or an AIN76-formula control diet for 3 weeks.

### Mucinase activity analysis

Mucinase activity was analyzed as described previously [79]. Briefly, fecal samples were mixed with acetate buffer (pH 5.5) at a ratio of 1:400 (w/v) and used for the mucinase assay with porcine stomach mucin as a substrate, according to a previously described method [80]. Fecal mucinase activity was expressed as the amount of liberated sugars per protein in the fecal homogenates per minute (nmol/mg protein/min).

### In vitro fermentation analysis

We employed the use of a previously described *in vitro* fermentation system [42]. Briefly, the cecal contents were obtained from rats fed an AIN93G diet for 7 days and diluted 50-fold in saline. Then, 110 mL of the diluted cecal contents was incubated in a jar fermenter at 37 ℃ under anaerobic, gently-stirred, and pH-controlled (pH > 5.2) conditions. After preincubation overnight, 3.3 g of porcine stomach mucin was added to the culture. To monitor the organic acid contents, 4 mL of each sample was collected 0, 24, 48 h after mucin supplementation.

### Preparation of colonic lamina propria cells

Colonic lymphocytes were prepared according to the methods of Weigmann *et al*. In brief, colonic tissues were treated with HBSS (Nakarai Tesque) containing 1 mM dithiothreitol and 20 mM EDTA (Nakarai Tesque) at 37 °C for 20 min to remove epithelial cells. The tissues were then minced and dissociated with RPMI 1640 medium (Nakarai Tesque) containing 0.5 mg/ml collagenase (Wako Pure Chemical Industries), 0.5 mg/mL DNase I (Merck, Darmstadt, Germany), 2 % FCS (MP Biomedicals, Santa Ana, CA), 100 U/mL penicillin, 100 μg/mL streptomycin, and 12.5 mM HEPES (pH 7.2) at 37 °C for 30 min to obtain a single-cell suspension which was filtered, washed with 2 % FCS in RPMI1640, and separated using a Percoll gradient.

### Flow cytometry

Cell staining was performed as described previously [81]. In brief, colonic lymphocytes were incubated with anti-mouse CD16/CD32 antibodies (93; BioLegend, Inc., San Diego, CA) to block their Fc receptors, then stained using antibodies conjugated with fluorescein isothiocyanate (FITC), phycoerythrin (PE), PerCP-Cyanine5.5, redFluor 710, eFluor 450, Brilliant Violet 510, or Brilliant Ultraviolet 737. Anti-CD45R/B220 (RA3-6B2), anti-IgA (C10-3), anti-RORγt (Q31-378), and anti-CD3e (145-2C11) antibodies were obtained from BD Biosciences (San Joses, CA); anti-CD3 (17A2) and anti-CD45R/B220 (RA3) antibodies were obtained from Tonbo Biosciences (San Diego, CA); anti-CD45 (30-F11) antibodies were obtained from BioLegend; and anti-Foxp3 (FJK-16s) and anti-CD4 (RM4-5) antibodies were obtained from Thermo Fisher Scientific (Waltham, MA). 7-AAD (BioLegend) was added to the cell suspension to label any dead cells. To visualize intracellular Foxp3 and RORγt, the cells were stained for surface antigens, fixed, and permeabilized using a Foxp3/Transcription Factor Staining Buffer Set (Themo Fisher Scientific). The permeabilized cells were then stained with anti-Foxp3 and anti-RORγt antibodies. Dead cells were detected using the Fixable Viability Stain 780 (BD Biosciences) for intracellular staining. The stained cells were analyzed using an LSR II Flow Cytometer (BD Biosciences) and FlowJo software ver. 10.4.2 (BD Biosciences).

### Statistical analysis

To analyze organic acid and mucin component levels, differences between two or more groups were analyzed using the Student’s *t*-test or ANOVA followed by Tukey’s multiple comparison test, respectively. When the variances were not homogeneous, the data were analyzed by the Wilcoxon’s rank sum test or the Kruskal-Wallis test followed by Dunn’s multiple comparison test. These statistical tests were performed in Prism ver. 7 (GraphPad Software, Inc., La Jolla, CA). The following statistical tests were performed in R 3.3.0. All correlation analyses were performed using Spearman’s correlation test with Benjamini–Hochberg false discovery rate (FDR) correction in the multtest R software package. Comparisons of bacterial taxon abundance were performed by LEfSe [38] using the default parameters, or the Kruskal-Wallis test followed by Dunn’s multiple comparison test in the dunn.test R software package. The biomarkers for each group identified using LEfSe were graphically annotated on to phylogenetic trees using GraPhlAn [82]. The correlation network was visualized using Cytoscape 3.3.0. No statistical methods were used to predetermine sample size. The experiments were not randomized. The investigators were not blinded to allocation during experiments and outcome assessment.

## Acknowledgements

We would like to thank Yun-Gi Kim for their comments and suggestions, and Koichiro Suzuki for their technical support.

## Author Contributions

K.H., T.M., J.K., and H.I. conceived the study. T.Y., Y.Fu., Y.Fr., and K.H. performed the microbiome analysis experiments. H.S and T.G analyzed mucin and metabolite levels and performed the rat experiments. R.N., H.H., and M.F. performed the *in vitro* fermentation analysis. T.Y., M.H., and Y.K. performed the mouse experiments. T.Y. and H.S. analyzed the data with M.H. and H.O. T.Y. performed the bioinformatic analysis with support from R.A., W.S. and M.H. H.I. collected clinical samples. T.Y., S.H., T.M., and K.H. interpreted the data. T.Y. and K.H. wrote the manuscript.

## Abbreviations

CD: Crohn’s disease
DNA: deoxyribonucleic acid
FDR: false discovery rate
HPLC: high performance liquid chromatography
IBD: inflammatory bowel disease
IgA: immunoglobulin A
OTU: operational taxonomic unit
PCR: polymerase chain reaction
RS: resistant starch
RNA: ribonucleic acid
rRNA: ribosomal RNA
SCFA: short chain fatty acid
Treg cell: regulatory T cell
Tris: tris(hydroxymethyl)aminomethane
UC: ulcerative colitis

## Supporting Information

**S1 Fig.**
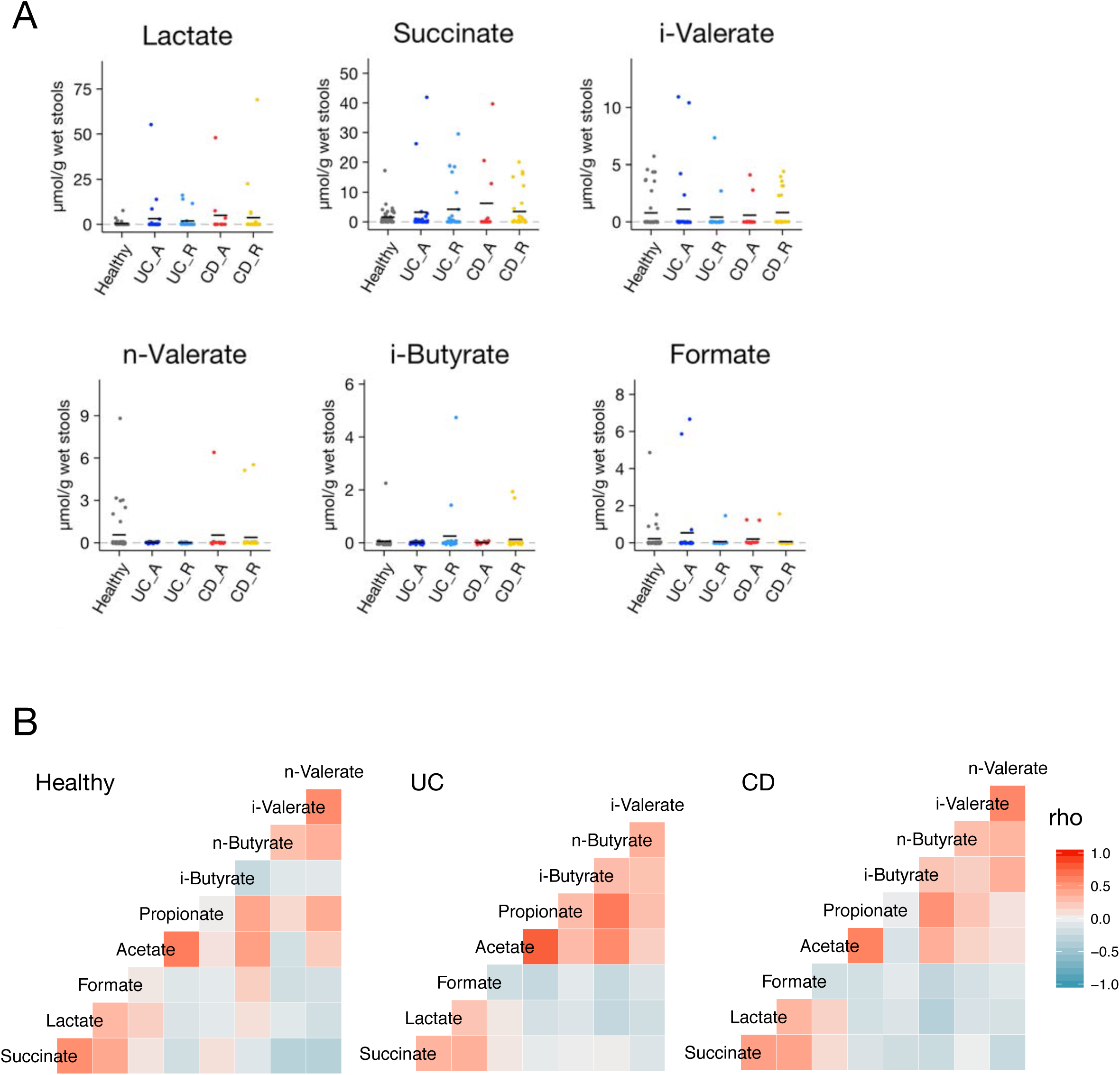
Organic acid profiles of the UC, CD, and healthy groups. (A) Fecal organic acid concentrations of the healthy subjects, patients with active UC (UC-A), remissive UC (UC-R), active CD (CD-A), and remissive CD (CD-R) were analyzed. Data represent the mean (*n* = 12-43/group). \**p* < 0.05 and \*\**p* < 0.01 (ANOVA followed by Tukey’s multiple comparison test or the Kruskal-Wallis test followed by Dunn’s multiple comparison test). (B) Heatmaps show the Spearman’s correlation coefficients of the fecal organic acid levels in each group, depicted as blue (negative correlation, −1) and red (positive correlation, +1) hues.

**S2 Fig.**
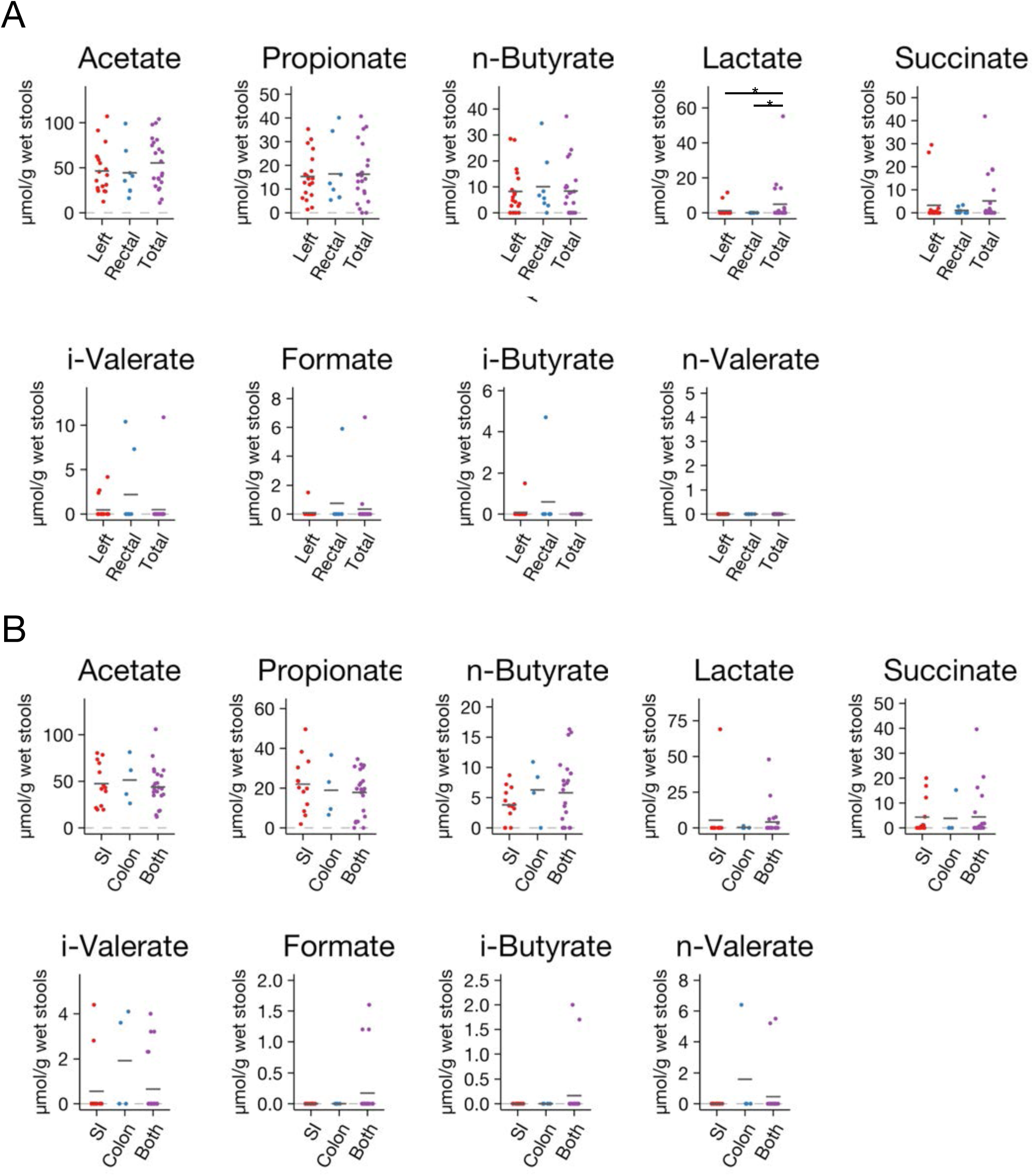
Association between fecal organic acids and lesion sites. Fecal organic acid concentrations of (A) UC patients or (B) CD patients grouped by lesion site were compared. Data represent the mean (*n* = 4-23/group). \**p* < 0.05 and \*\**p* < 0.01 (ANOVA followed by Tukey’s multiple comparison test or the Kruskal-Wallis test followed by Dunn’s multiple comparison test).

**S3 Fig.**
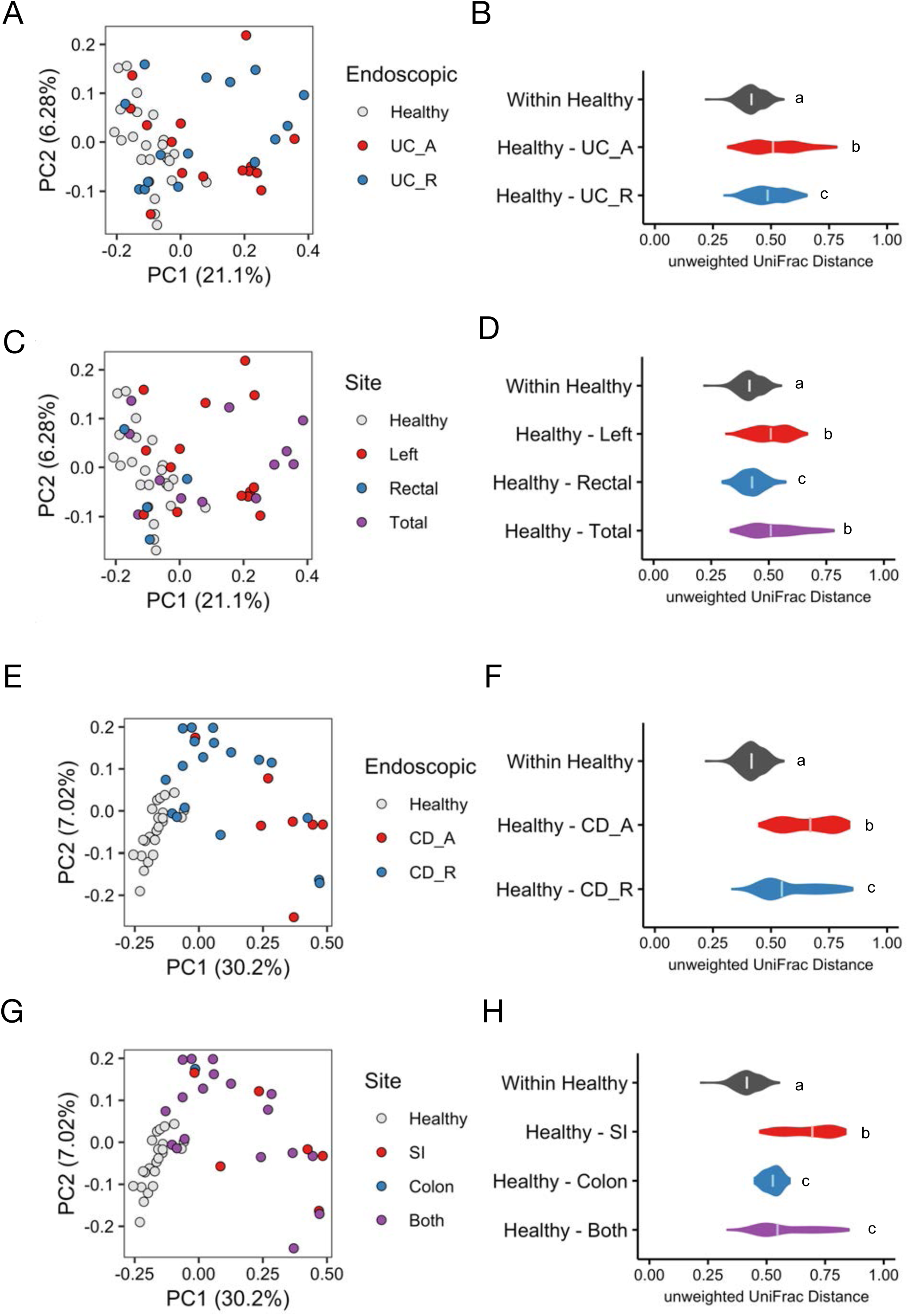
Association between fecal bacterial composition and endoscopic assessment or lesion site. (A, C, E, and G) Principal coordinate analysis of the unweighted UniFrac distance between the microbiota of UC patients grouped by (A) endoscopic assessment or (B) lesion site, and CD patients grouped by (E) endoscopic assessment or (G) lesion sites. (B, D, F, and H) Violin plots of the unweighted UniFrac distance between the microbiota of the healthy subjects. Median values are shown. Statistical analysis was performed using the Kruskal-Wallis test followed by Dunn’s multiple comparison test (n = 7-23/group). The letters over the boxplots indicate significant differences (*p* < 0.05).

**S4 Fig.**
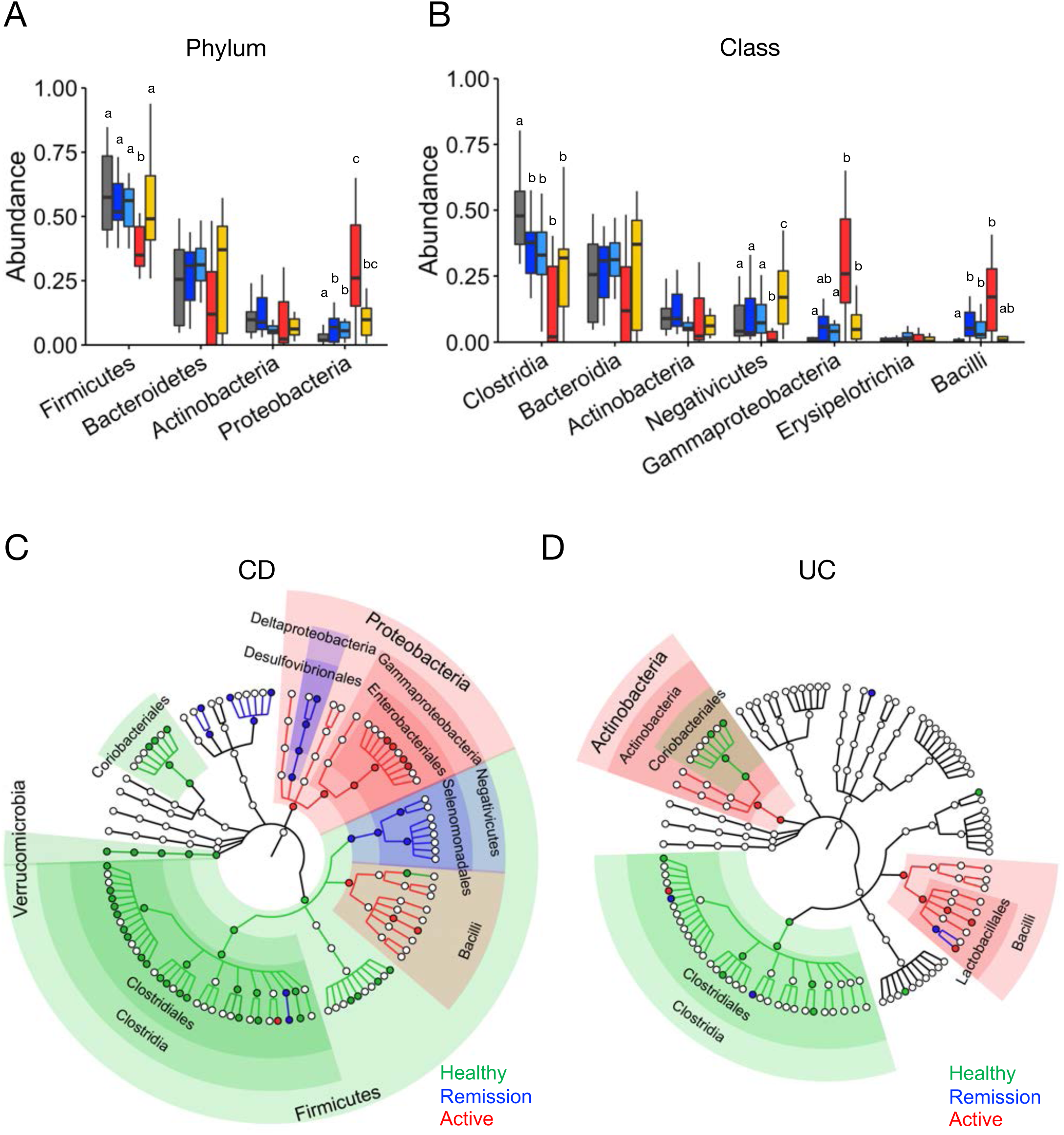
Microbial community comparison between IBD patients and healthy subjects. Boxplots of relative bacterial abundance at the phylum (A) and class (B) levels. Each boxplot represents the median, IQR, and the lowest and highest values within 1.5 IQRs of the first and third quartiles. The outliers are not shown. Statistical analysis was performed using the Kruskal-Wallis test followed by Dunn’s multiple comparison test (n = 7-23/group). The letters over the boxplots indicate significant differences (*p* < 0.05). (C, D) Taxa discriminating between microbiota of (C) UC or (D) CD patients in active or remission phase and healthy subjects, as determined by LEfSe [38] analysis. The discriminative taxa are annotated on phylogenetic trees.

**S5 Fig.**
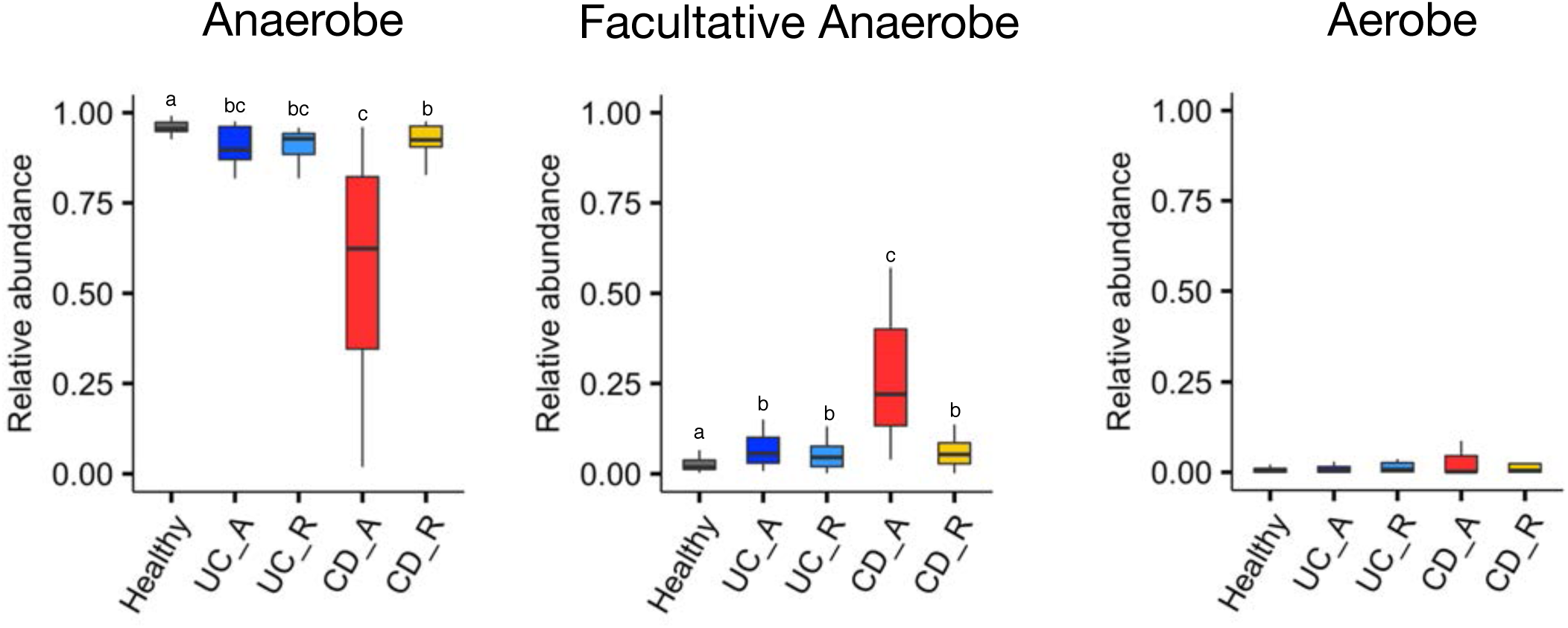
Occupancy of anaerobes, facultative anaerobes, and aerobes in the microbial communities of IBD patients and healthy subjects. Relative abundance of anaerobic, facultatively anaerobic, and aerobic bacteria was predicted using BugBase. Each boxplot represents the median, IQR, and the lowest and highest values within 1.5 IQRs of the first and third quartiles. The outliers are not shown. Statistical analysis was performed using the Kruskal-Wallis test followed by Dunn’s multiple comparison test (n = 7-23/group). The letters over the boxplots indicate significant differences (*p* < 0.05).

**S6 Fig.**
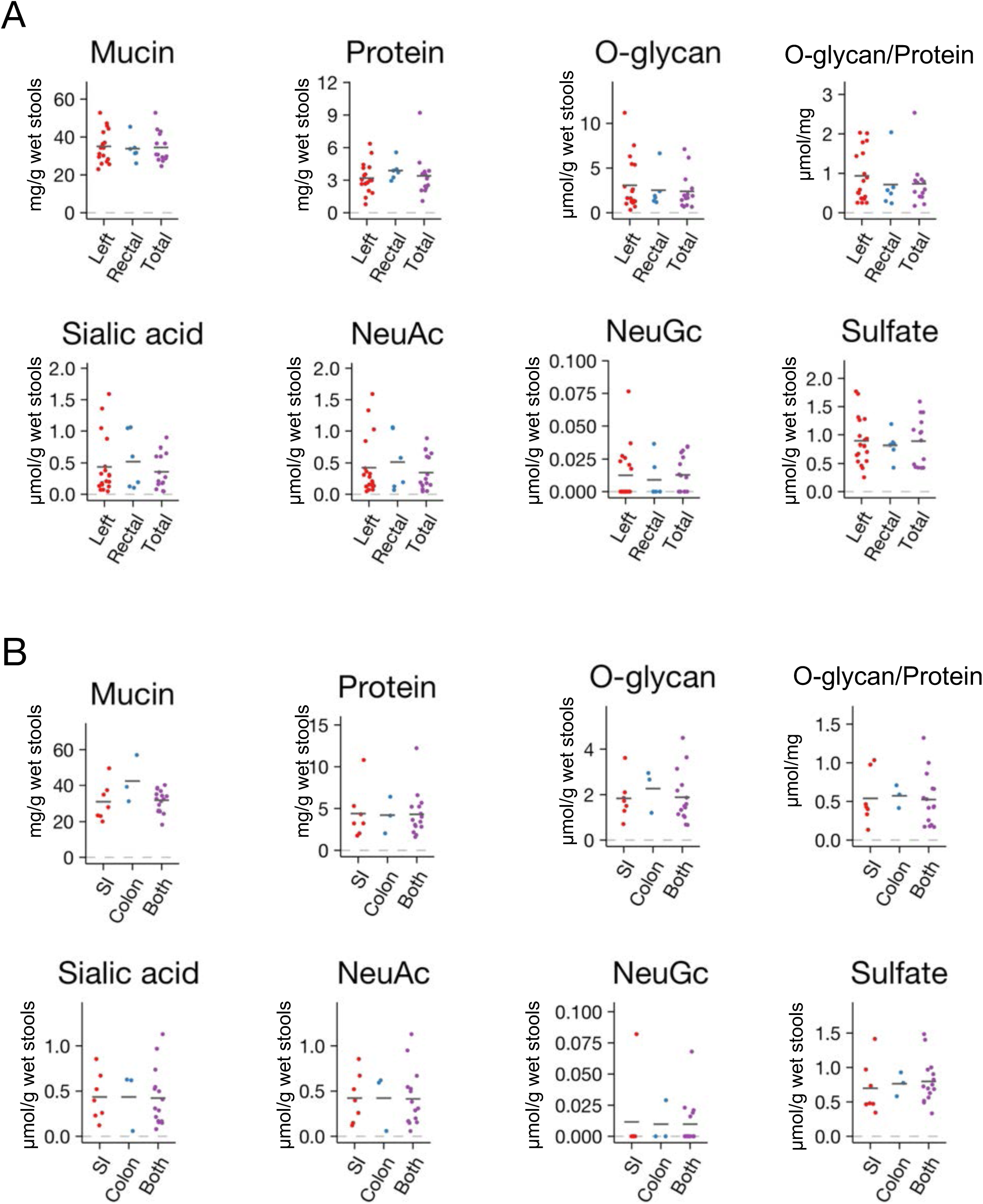
Association between stool mucin components and lesion sites. Stool mucin component concentrations and mucin *O*-glycan ratios per protein were compared between the (A) UC patients and (B) CD patients grouped by lesion site. Data represent the mean (*n* = 3-18/group). \**p* < 0.05 and \*\**p* < 0.01 (ANOVA followed by Tukey’s multiple comparison test or the Kruskal-Wallis test followed by Dunn’s multiple comparison test).

**S7 Fig.**
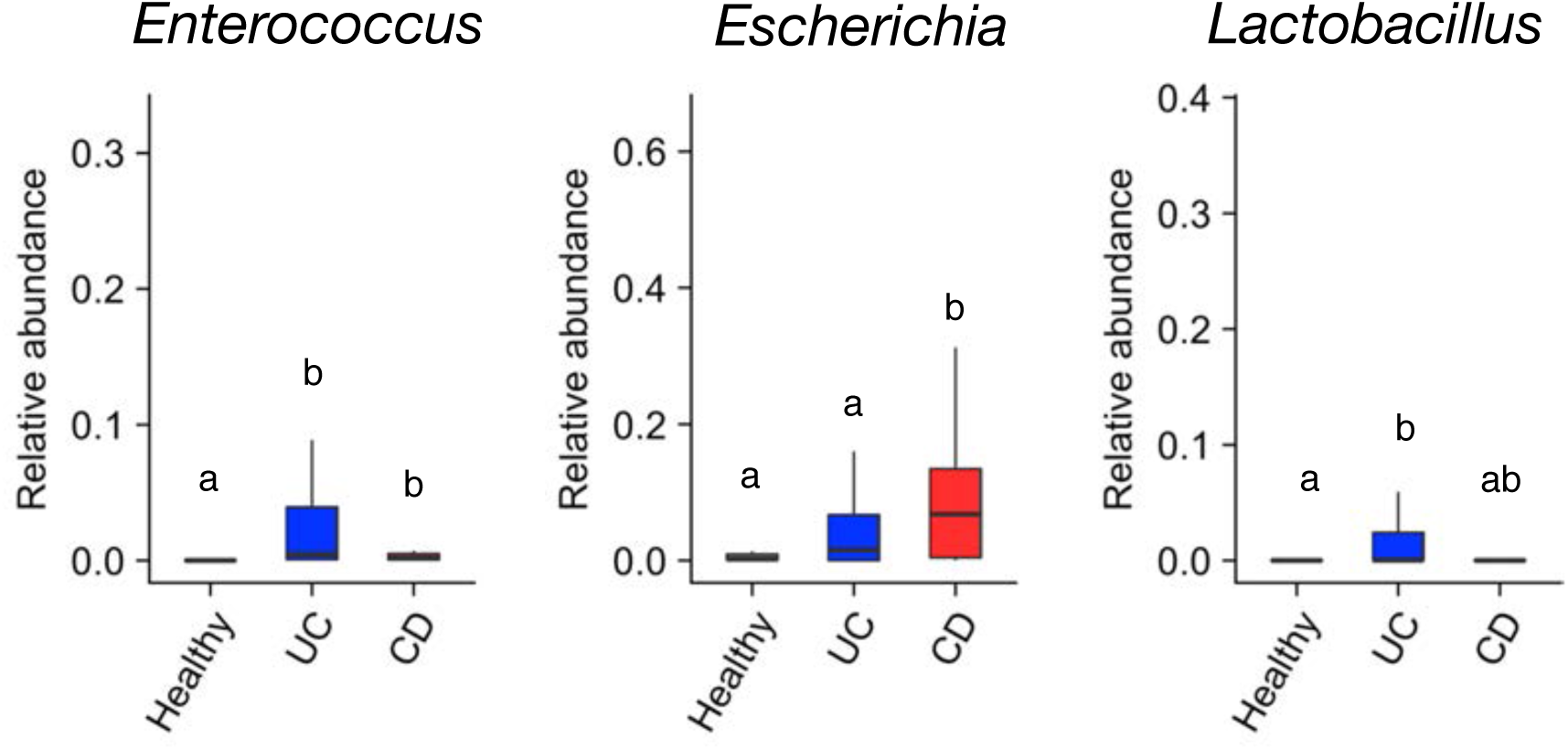
Bacterial genera overrepresented in IBD patients. Boxplots of the relative abundances of bacterial genera. Each boxplot represents the median, IQR, and the lowest and highest values within 1.5 IQRs of the first and third quartiles. The outliers are not shown. Statistical analysis was performed using the Kruskal-Wallis test followed by Dunn’s multiple comparison test (n = 7-23/group). The letters over the boxplots indicate significant differences (*p* < 0.05).

**S8 Fig.**
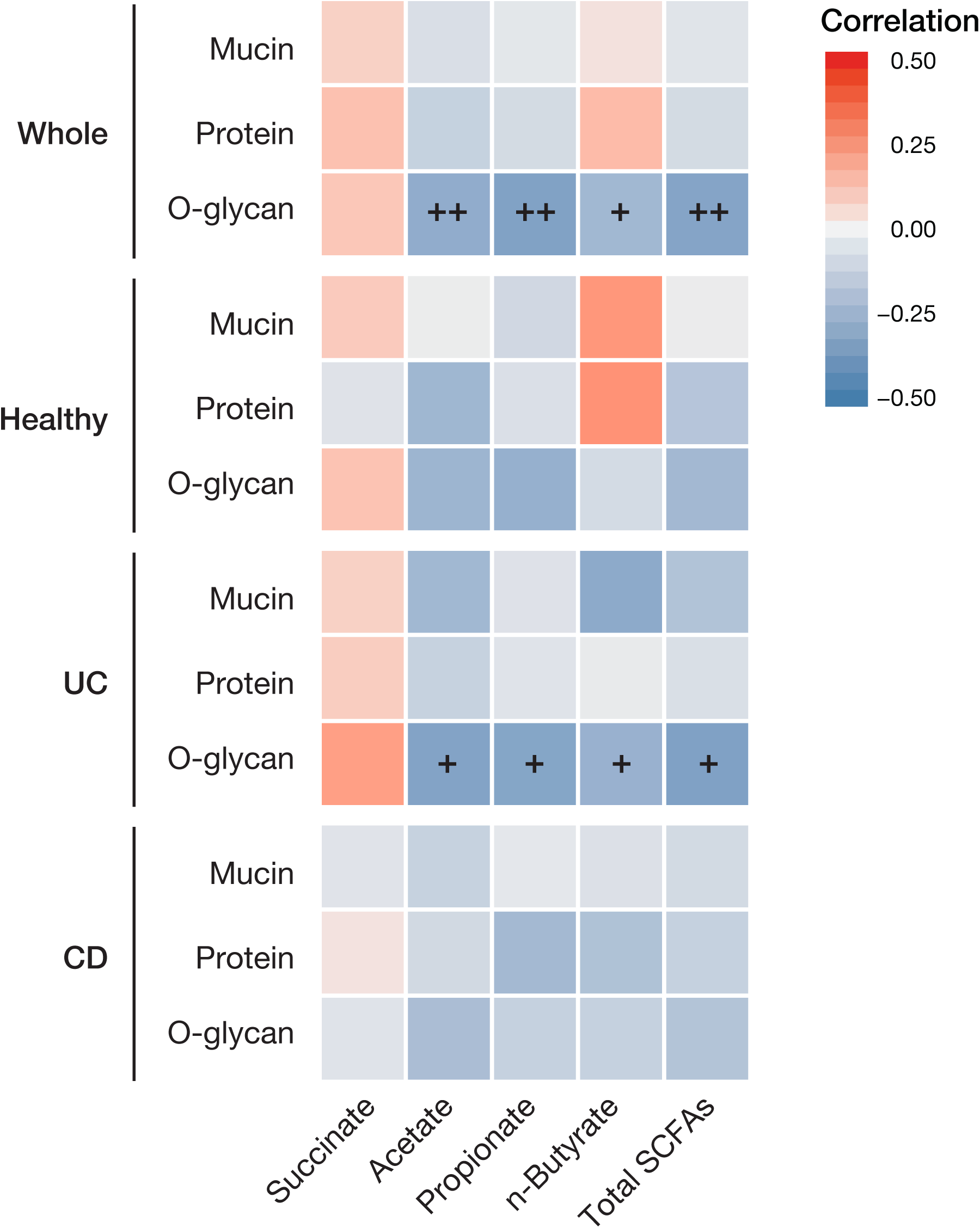
Correlation between the fecal SCFAs and mucin components of the individual disease groups. Correlation analysis between the concentrations of major stool organic acids and mucin components in the stool samples. Heatmaps of the Spearman’s correlation coefficients, depicted as blue (negative correlation, −0.5) and red (positive correlation, +0.5) hues. The Benjamini-Hochberg (BH) method was used for FDR adjustment in each group. +: FDR < 0.05, ++: FDR < 0.01.

**S9 Fig.**
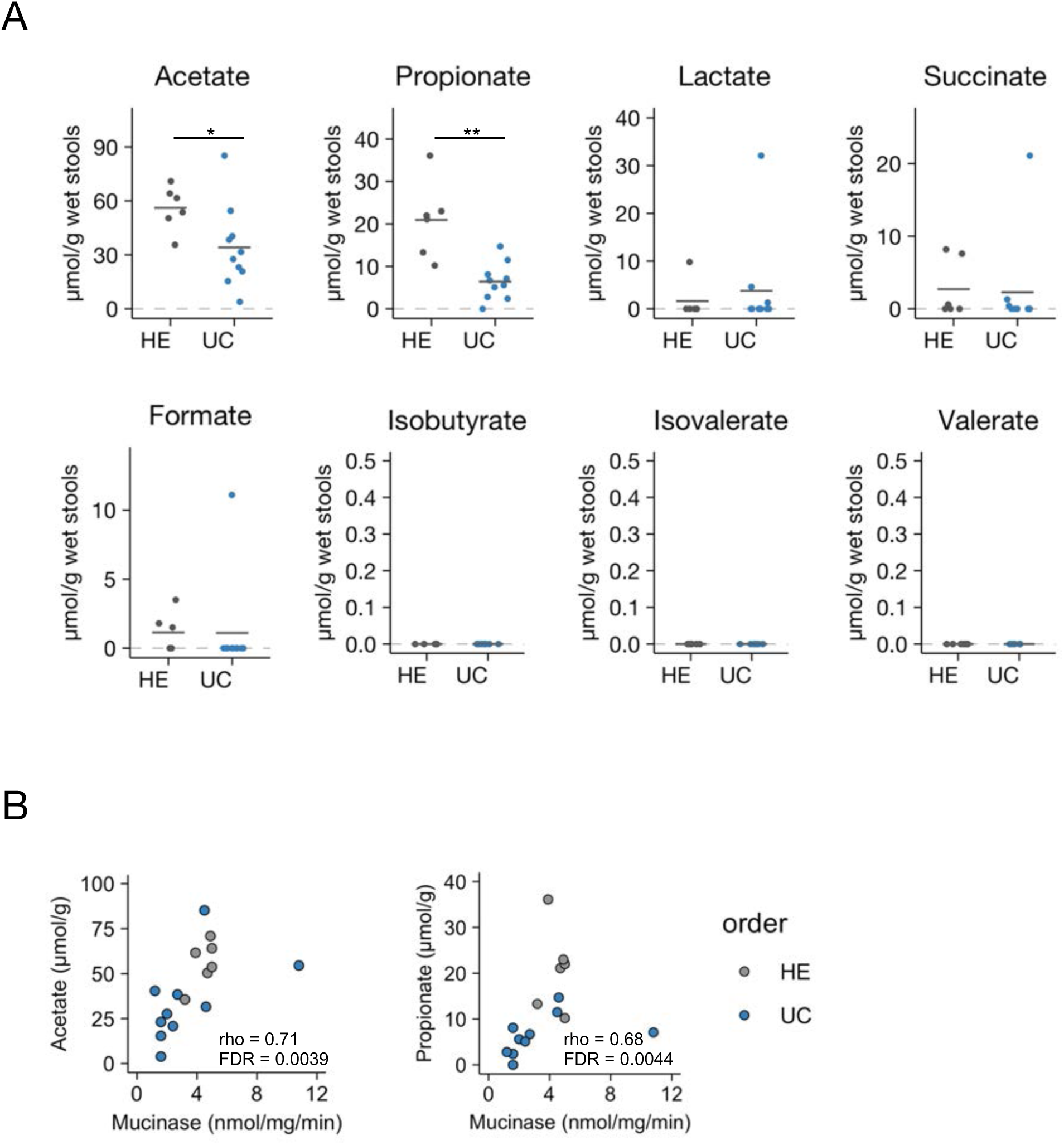
Association between mucinase activity and fecal SCFA levels in UC patients and healthy subjects. (A) Organic acid concentrations were analyzed in the stool samples of healthy subjects and UC patients. Data represent the mean (*n* = 6, 10/group). \**p* < 0.05 and \*\**p* < 0.01 (Student’s *t* test or Wilcoxon’s rank sum test). (B) Scatter plots of mucinase activity versus the concentrations of acetate or propionate. The Spearman’s correlation coefficient (rho) and FDR are shown.

**S10 Fig.**
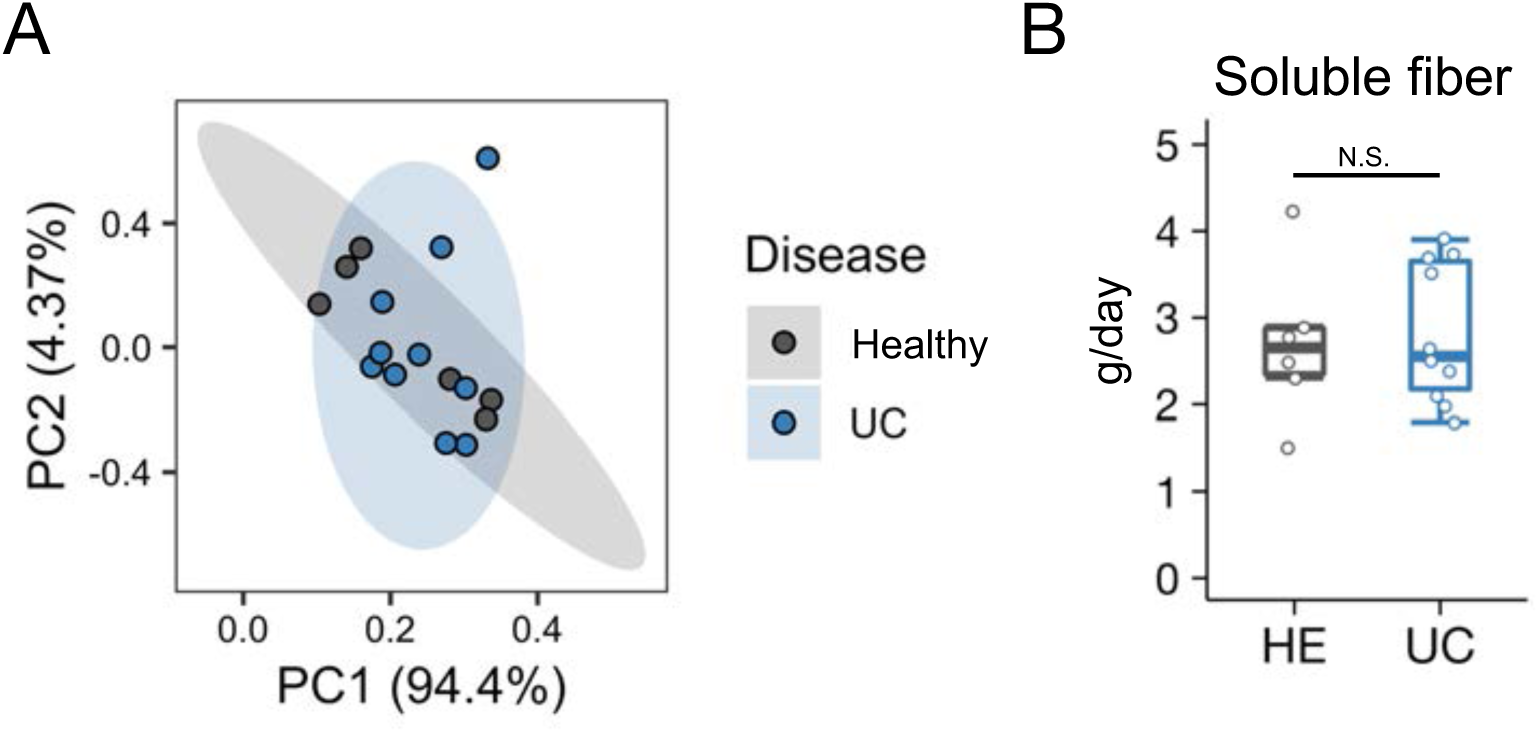
Comparison of nutrient intake between UC patients and healthy subjects. (A) Principal component analysis plot of the nutrient intake data in S9 Table. Standard error (95 %) ellipses are shown around the UC patients and healthy subjects. (B) Boxplot of soluble fiber intake indicating the median, IQR, and the lowest and highest values within 1.5 IQRs of the first and third quartiles. NS, not statistically significant (Student’s *t* test, *n* = 6, 10).

**S11 Fig.**
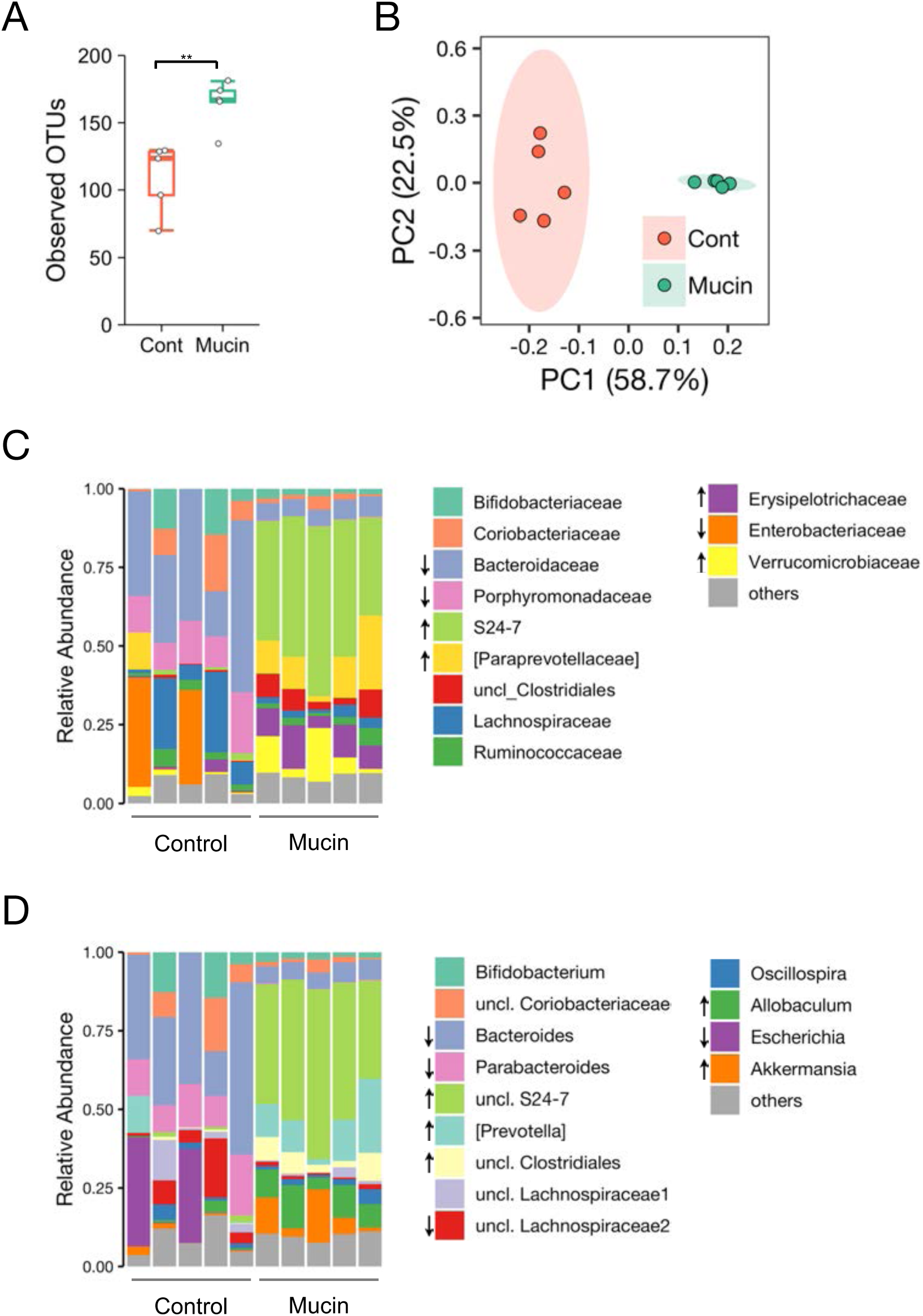
Effects of mucin administration on cecal microbiota in mice. (A) Species richness (number of observed OTUs) of the microbiota of control or mucin-fed mice.\**p* < 0.05 and \*\**p* < 0.01 (Wilcoxon’s rank sum test). (B) Principal coordinate analysis of the weighted UniFrac distance between the control and mucin-fed mice. (C, D) Staked bar plot of bacterial abundances at family (C) and genus (D) levels. Statistical analysis was performed using LEfSe (*n* = 5/group).

**S12 Fig.**
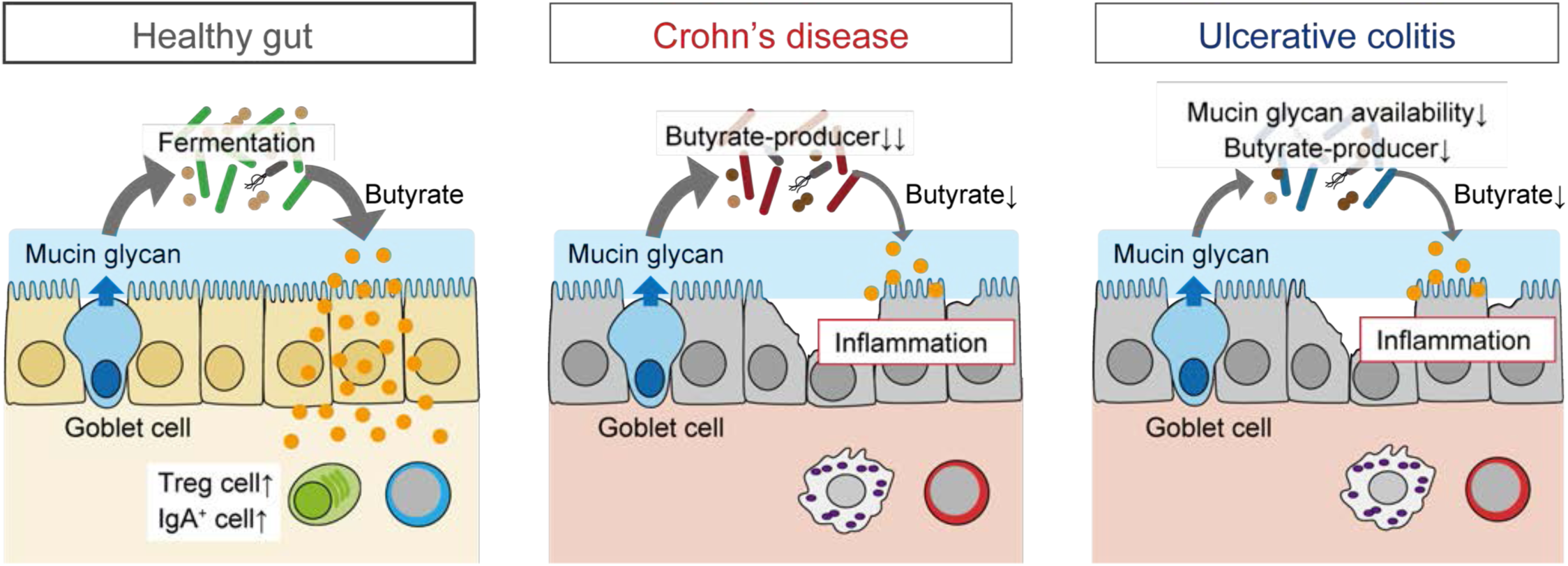
Distinct dysbiosis models in CD and UC. In the healthy gut, intestinal microbiota utilize mucin *O*-glycan as an endogenous fermentation source to produce butyrate (left). In Crohn’s disease, major butyrate producers such as *Faecalibacterium prausnitzii* are almost absent due to severe dysbiosis, resulting in the loss of luminal butyrate (center). In contrast, the gut microbiota of UC patients showed less severe dysbiosis and normal levels of *F. prausnitzii*. However, *O*-glycan availability was compromised in the UC-associated microbiota, leading to a substantial decrease in luminal butyrate (right). Reduced butyrate production may render the hosts susceptible to chronic inflammation in the gut.

**S1 Table.** UC patient information (Study 1). The number and ratio of patients with different lesion sites, disease severity score, or drug treatments.

| Ulcerative colitis (n=49) |  |
| --- | --- |
| Site of disease |  |
| Total (%) | 22 (44.9%) |
| Rectal (%) | 8 (16.3%) |
| Left (%) | 19 (38.8%) |
| Matts Endoscopy Score |  |
| 1 | 24 (49.0%) |
| 2 | 14 (28.6%) |
| 3 | 6 (12.2%) |
| 4 | 5 (10.2%) |
| Clinical Activity Index |  |
| 0 – 3 | 38 (22.4%) |
| 4 ≤ | 11 (77.6%) |
| UC Drugs |  |
| Anti-TNF antibody | 1 (2.0%) |
| Immunosuppressant | 11 (22.4%) |
| Prodrug of 5-ASA | 34 (69.4%) |
| PPI | 7 (14.3%) |
| Steroid | 7 (14.3%) |
| Probiotics | 40 (81.6%) |

**S2 Table.** CD patient information (Study 1). The number and ratio of patients with different lesion sites, disease severity score, or drug treatments.

| Crohn's Disease (n = 40) |  |
| --- | --- |
| Site of disease |  |
| Small Intestine (%) | 13 (32.5 %) |
| Colon (%) | 4 (10 %) |
| Both (%) | 23 (57.5 %) |
| Modified Rutgeerts score |  |
| 0-1 (remission) | 12 (30 %) |
| 2-4 (active) | 28 (70 %) |
| CDAI |  |
| <150 | 30 (75 %) |
| 150 – 220 | 6 (15 %) |
| 220 – 300 | 2 (5 %) |
| 300< | 2 (5 %) |
| CD Drugs |  |
| Anti-TNF antibody | 23 (57.5 %) |
| Immunosuppressant | 13 (32.5 %) |
| Prodrug of 5-ASA | 34 (85.0%) |
| PPI | 1 (2.5 %) |
| Antibiotics | 2 (5.0 %) |
| Probiotics | 27 (67.5 %) |

**S3 Table.** Effects of lesion site and drug treatment on fecal *n*-butyrate levels in UC patients (Study 1). Data represents the mean ± SD of fecal *n*-butyrate levels (μmol/g) and the *p* value of UC patients with different lesion sites or drug treatments.

|  | Total | Left | Rectal | *p* value |
| --- | --- | --- | --- | --- |
| Site of disease | 8.42±10.0 | 8.28±8.65 | 10.2±11.4 | 0.89 (ANOVA) |
|  | Treated | Untreated | *p* value |  |
| Immunosuppressant | 5.73±8.06 | 9.48±9.87 | 0.26 (Student's t test) |  |
| Prodrug of 5-ASA | 8.21±8.89 | 12.44±14.96 | 0.35 (Student's t test) |  |
| PPI | 10.83±9.62 | 8.27±9.60 | 0.52 (Student's t test) |  |
| Steroid | 4.96±10.49 | 9.25±9.37 | 0.28 (Student's t test) |  |
| Probiotics | 8.39±10.12 | 10.51±5.89 | 0.71 (Student's t test) |  |

**S4 Table.** Effects of lesion site and drug treatment on fecal *n*-butyrate levels in CD patients (Study 1). Data represents the mean ± SD of fecal *n*-butyrate levels (μmol/g) and the *p* value of CD patients with different lesion sites or drug treatments.

|  | Small Intestine | Colon | Both | *p* value |
| --- | --- | --- | --- | --- |
| Site of disease | 8.42±10.0 | 8.28±8.65 | 10.2±11.4 | 0.62 (Kruskal-Wallis) |
|  | Treated | Untreated | *p* value |  |
| Anti-TNF | 5.30±4.84 | 5.11±4.52 | 0.90 (Student's t test) |  |
| Immunosuppressant | 6.04±5.08 | 4.82±4.47 | 0.45 (Student's t test) |  |
| Prodrug of 5-ASA | 5.26±4.52 | 4.98±5.76 | 0.90 (Student's t test) |  |
| PPI | 2.80±1.11 | 5.41±4.77 | 0.52 (Student's t test) |  |
| Probiotics | 5.70±5.20 | 4.22±3.17 | 0.35 (Student's t test) |  |

**S5 Table.** Summary of bacterial genera associated with fecal organic acid levels (Study 1). Data represent the Spearman’s correlation coefficients, *p* values with Benjamini-Hochberg correction (FDR), and average bacterial abundance of healthy subjects (%). Bacteria abundant (more than 1 %) in the healthy subjects are shown in bold.

n-Butyrate
| Bacterial genera | rho | FDR | Abundance in healthy subjects (%) |
| --- | --- | --- | --- |
| *Haemophilus* | 0.55 | 0.000028 | 0.18 |
| ***Faecalibacterium*** | 0.53 | 0.000029 | 11.81 |
| ***Eubacterium*** | 0.51 | 0.000056 | 7.62 |
| *Roseburia* | 0.45 | 0.0010 | 0.79 |
| *Coprobacter* | 0.42 | 0.0031 | 0.03 |
| ***Anaerostipes*** | 0.39 | 0.0066 | 1.79 |
| *Eggerthella* | 0.35 | 0.022 | 0.33 |
| *Turicibacter* | 0.34 | 0.029 | 0.05 |
| *Parasutterella* | 0.34 | 0.029 | 0.39 |
| *Dorea* | 0.33 | 0.029 | 0.63 |
| ***Ruminococcus*** | 0.32 | 0.041 | 4.16 |
| ***Gemmiger*** | 0.32 | 0.041 | 2.67 |

Acetate
| Bacterial genera | rho | FDR | Abundance in healthy subjects (%) |
| --- | --- | --- | --- |
| *Proteus* | 0.46 | 0.0022 | 0.0017 |
| *Coprobacter* | 0.44 | 0.0022 | 0.03 |
| *Haemophilus* | 0.44 | 0.0022 | 0.18 |

n-Valerate
| Bacterial genera | rho | FDR | Abundance in healthy subjects (%) |
| --- | --- | --- | --- |
| *Faecalitalea* | 0.39 | 0.042 | 0.026 |

i-Butyrate
| Bacterial genera | rho | FDR | Abundance in healthy subjects (%) |
| --- | --- | --- | --- |
| *Bacillus* | 0.39 | 0.049 | 0.0078 |

**S6 Table.** Summary of bacterial species associated with fecal organic acid levels (Study 1). Data represent the Spearman’s correlation coefficients, *p* values with Benjamini-Hochberg correction (FDR), and average bacterial abundance of healthy subjects (%). Bacteria abundant (more than 1 %) in the healthy subjects are shown in bold.

n-Butyrate
| Bacterial species | rho | FDR | Abundance in healthy subjects (%) |
| --- | --- | --- | --- |
| *Haemophilus parainfluenzae* | 0.55 | 0.000037 | 0.18 |
| *Eubacterium ventriosum* | 0.54 | 0.000037 | 0.60 |
| ***Faecalibacterium prausnitzii*** | 0.53 | 0.000044 | 11.64 |
| *Ruminococcus obeum* | 0.48 | 0.00054 | 0.58 |
| ***Eubacterium rectale*** | 0.48 | 0.00056 | 2.38 |
| *Faecalibacterium sp.* | 0.47 | 0.00060 | 0.08 |
| *Eubacterium ramulus* | 0.46 | 0.00084 | 0.29 |
| ***Eubacterium hallii*** | 0.43 | 0.0023 | 2.04 |
| *Anaerostipes hadrus* | 0.42 | 0.0029 | 0.48 |
| *Roseburia inulinivorans* | 0.42 | 0.0030 | 0.21 |
| *Coprobacter fastidiosus* | 0.42 | 0.0032 | 0.03 |
| *Blautia glucerasea* | 0.39 | 0.0067 | 0.25 |
| ***Anaerostipes sp.*** | 0.39 | 0.0067 | 1.28 |
| *Roseburia intestinalis* | 0.39 | 0.0067 | 0.43 |
| *Roseburia faecis* | 0.37 | 0.014 | 0.04 |
| *Haemophilus pittmaniae* | 0.36 | 0.017 | 0.01 |
| *Eggerthella lenta* | 0.35 | 0.020 | 0.33 |
| *Phascolarctobacterium sp.* | 0.35 | 0.020 | 0.25 |
| *Ruminococcus gauvreauii* | 0.33 | 0.027 | 0.01 |
| *Clostridium xylanolyticum* | 0.34 | 0.027 | 0.11 |
| *Parasutterella excrementihominis* | 0.34 | 0.027 | 0.39 |
| *Ruminococcus faecis* | 0.34 | 0.027 | 0.89 |
| *Turicibacter sanguinis* | 0.34 | 0.027 | 0.05 |
| *Clostridium disporicum* | 0.33 | 0.028 | 0.05 |
| *Blautia schinkii* | 0.32 | 0.033 | 0.19 |
| ***Ruminococcus bromii*** | 0.31 | 0.043 | 2.12 |
| ***Blautia wexlerae*** | 0.31 | 0.045 | 3.16 |
| ***Bacteroides vulgatus*** | 0.31 | 0.049 | 4.05 |

Acetate
| Bacterial species | rho | FDR | Abundance in healthy subjects (%) |
| --- | --- | --- | --- |
| *Coprobacter fastidiosus* | 0.44 | 0.0089 | 0.030 |
| *Harmophilus parainfluenzae* | 0.43 | 0.0089 | 0.17 |

n-Valerate
| Bacterial species | rho | FDR | Abundance in healthy subjects (%) |
| --- | --- | --- | --- |
| *Ruminococcus gauvreauii* | 0.52 | 0.00035 | 0.013 |
| *Eubacterium cylindroides* | 0.39 | 0.048 | 0.026 |

**S7 Table.**
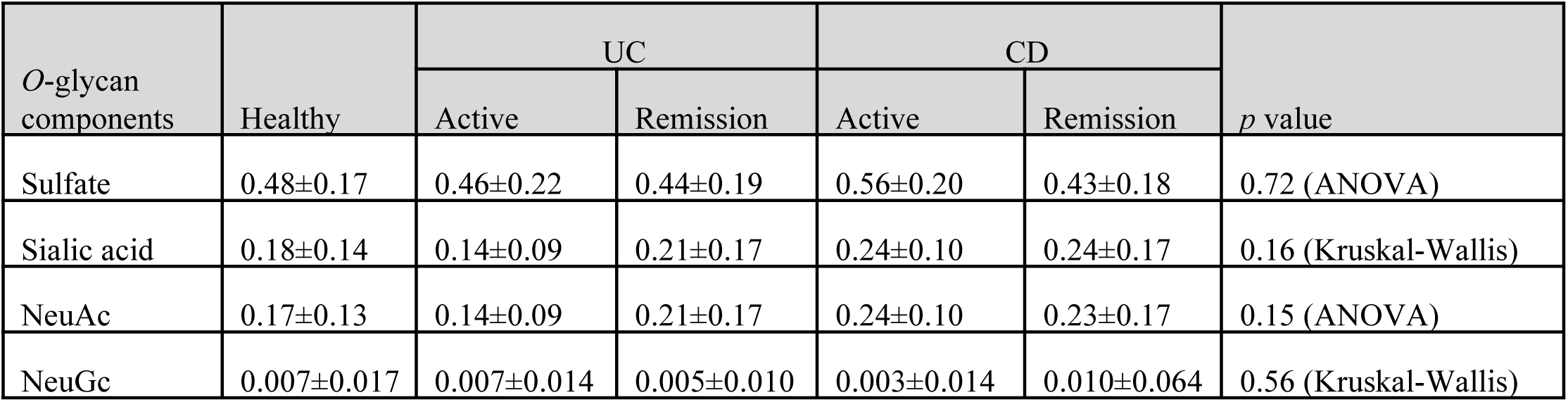
Association between IBD and stool mucin *O*-glycan component ratio (Study 1). Data represent the mean ± SD of the ratio of each mucin *O*-glycan component per mucin *O*-glycan (nmol/nmol) and the *p* values of the healthy subjects and IBD patients classified by endoscopic assessment.

| *O*-glycan components | Healthy | UC |  | CD |  | *p* value |
| --- | --- | --- | --- | --- | --- | --- |
|  |  | Active | Remission | Active | Remission |  |
| Sulfate | 0.48±0.17 | 0.46±0.22 | 0.44±0.19 | 0.56±0.20 | 0.43±0.18 | 0.72 (ANOVA) |
| Sialic acid | 0.18±0.14 | 0.14±0.09 | 0.21±0.17 | 0.24±0.10 | 0.24±0.17 | 0.16 (Kruskal-Wallis) |
| NeuAc | 0.17±0.13 | 0.14±0.09 | 0.21±0.17 | 0.24±0.10 | 0.23±0.17 | 0.15 (ANOVA) |
| NeuGc | 0.007±0.017 | 0.007±0.014 | 0.005±0.010 | 0.003±0.014 | 0.010±0.064 | 0.56 (Kruskal-Wallis) |

**S8 Table.** Association between IBD and fecal NeuGc detection frequency (Study 1). Data represent the proportion of subjects in whom NeuGc was detected. Statistical tests were performed using Fisher’s exact test by comparing with healthy subjects.

| Disease | Detection ratio | *p* value |
| --- | --- | --- |
| Healthy | 8/38 (21%) | - |
| UC | 17/38 (45%) | 0.0497 |
| CD | 7/25 (28%) | 0.558 |

**S9 Table.**
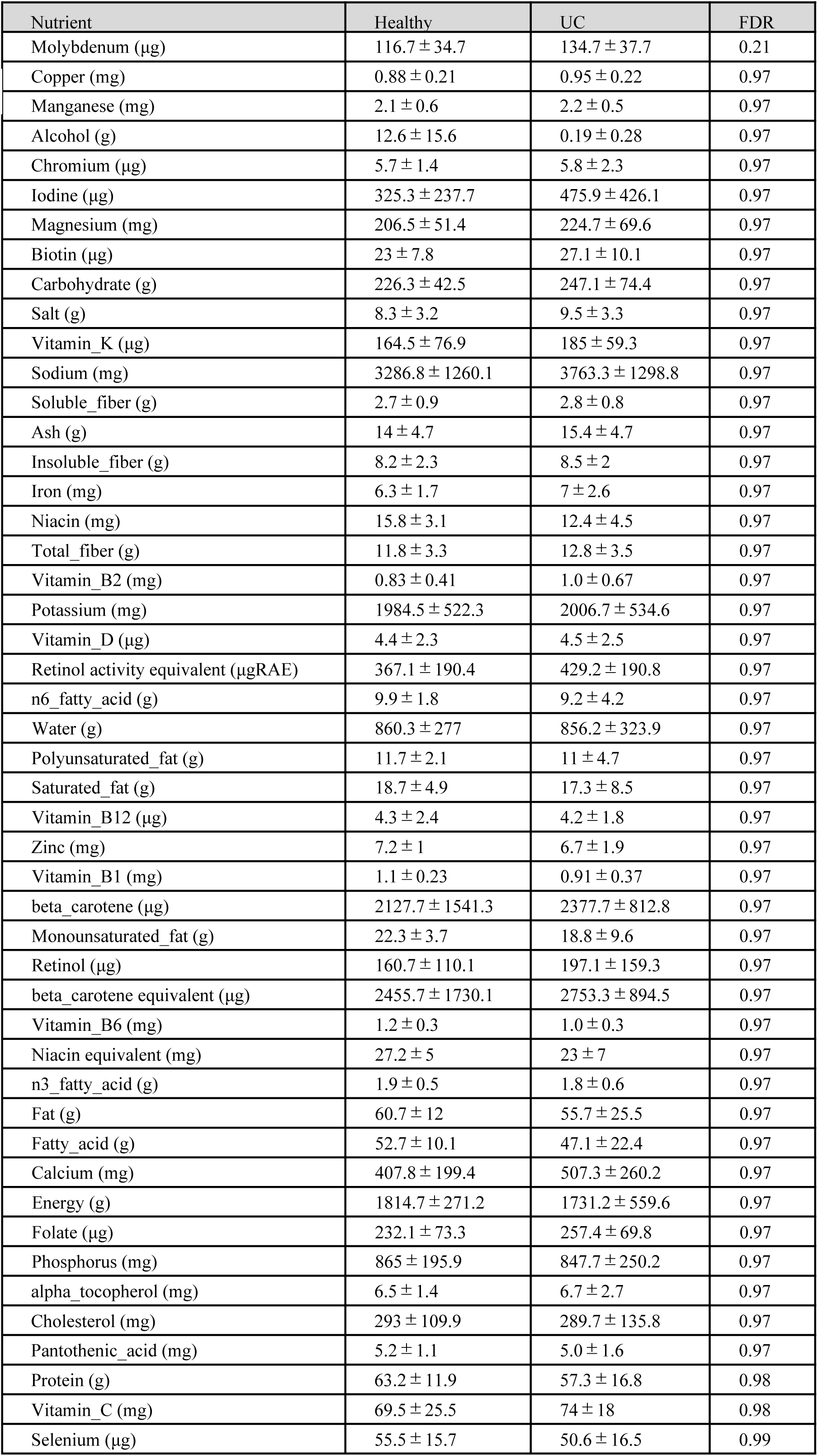
Summary of the nutrient intake of healthy subjects and UC patients (Study 2). Data represent the mean ± SD of the daily nutrient intake and *p* values with Benjamini-Hochberg correction (FDR).

| Nutrient | Healthy | UC | FDR |
| --- | --- | --- | --- |
| Molybdenum (µg) | 116.7 ± 34.7 | 134.7 ± 37.7 | 0.21 |
| Copper (mg) | 0.88 ± 0.21 | 0.95 ± 0.22 | 0.97 |
| Manganese (mg) | 2.1 ± 0.6 | 2.2 ± 0.5 | 0.97 |
| Alcohol (g) | 12.6 ± 15.6 | 0.19 ± 0.28 | 0.97 |
| Chromium (µg) | 5.7 ± 1.4 | 5.8 ± 2.3 | 0.97 |
| Iodine (µg) | 325.3 ± 237.7 | 475.9 ± 426.1 | 0.97 |
| Magnesium (mg) | 206.5 ± 51.4 | 224.7 ± 69.6 | 0.97 |
| Biotin (µg) | 23 ± 7.8 | 27.1 ± 10.1 | 0.97 |
| Carbohydrate (g) | 226.3 ± 42.5 | 247.1 ± 74.4 | 0.97 |
| Salt (g) | 8.3 ± 3.2 | 9.5 ± 3.3 | 0.97 |
| Vitamin_K (µg) | 164.5 ± 76.9 | 185 ± 59.3 | 0.97 |
| Sodium (mg) | 3286.8 ± 1260.1 | 3763.3 ± 1298.8 | 0.97 |
| Soluble_fiber (g) | 2.7 ± 0.9 | 2.8 ± 0.8 | 0.97 |
| Ash (g) | 14 ± 4.7 | 15.4 ± 4.7 | 0.97 |
| Insoluble_fiber (g) | 8.2 ± 2.3 | 8.5 ± 2 | 0.97 |
| Iron (mg) | 6.3 ± 1.7 | 7 ± 2.6 | 0.97 |
| Niacin (mg) | 15.8 ± 3.1 | 12.4 ± 4.5 | 0.97 |
| Total_fiber (g) | 11.8 ± 3.3 | 12.8 ± 3.5 | 0.97 |
| Vitamin_B2 (mg) | 0.83 ± 0.41 | 1.0 ± 0.67 | 0.97 |
| Potassium (mg) | 1984.5 ± 522.3 | 2006.7 ± 534.6 | 0.97 |
| Vitamin_D (µg) | 4.4 ± 2.3 | 4.5 ± 2.5 | 0.97 |
| Retinol activity equivalent (µgRAE) | 367.1 ± 190.4 | 429.2 ± 190.8 | 0.97 |
| n6_fatty_acid (g) | 9.9 ± 1.8 | 9.2 ± 4.2 | 0.97 |
| Water (g) | 860.3 ± 277 | 856.2 ± 323.9 | 0.97 |
| Polyunsaturated_fat (g) | 11.7 ± 2.1 | 11 ± 4.7 | 0.97 |
| Saturated_fat (g) | 18.7 ± 4.9 | 17.3 ± 8.5 | 0.97 |
| Vitamin_B12 (µg) | 4.3 ± 2.4 | 4.2 ± 1.8 | 0.97 |
| Zinc (mg) | 7.2 ± 1 | 6.7 ± 1.9 | 0.97 |
| Vitamin_B1 (mg) | 1.1 ± 0.23 | 0.91 ± 0.37 | 0.97 |
| beta_carotene (µg) | 2127.7 ± 1541.3 | 2377.7 ± 812.8 | 0.97 |
| Monounsaturated_fat (g) | 22.3 ± 3.7 | 18.8 ± 9.6 | 0.97 |
| Retinol (µg) | 160.7 ± 110.1 | 197.1 ± 159.3 | 0.97 |
| beta_carotene equivalent (µg) | 2455.7 ± 1730.1 | 2753.3 ± 894.5 | 0.97 |
| Vitamin_B6 (mg) | 1.2 ± 0.3 | 1.0 ± 0.3 | 0.97 |
| Niacin equivalent (mg) | 27.2 ± 5 | 23 ± 7 | 0.97 |
| n3_fatty_acid (g) | 1.9 ± 0.5 | 1.8 ± 0.6 | 0.97 |
| Fat (g) | 60.7 ± 12 | 55.7 ± 25.5 | 0.97 |
| Fatty_acid (g) | 52.7 ± 10.1 | 47.1 ± 22.4 | 0.97 |
| Calcium (mg) | 407.8 ± 199.4 | 507.3 ± 260.2 | 0.97 |
| Energy (g) | 1814.7 ± 271.2 | 1731.2 ± 559.6 | 0.97 |
| Folate (µg) | 232.1 ± 73.3 | 257.4 ± 69.8 | 0.97 |
| Phosphorus (mg) | 865 ± 195.9 | 847.7 ± 250.2 | 0.97 |
| alpha_tocopherol (mg) | 6.5 ± 1.4 | 6.7 ± 2.7 | 0.97 |
| Cholesterol (mg) | 293 ± 109.9 | 289.7 ± 135.8 | 0.97 |
| Pantothenic_acid (mg) | 5.2 ± 1.1 | 5.0 ± 1.6 | 0.97 |
| Protein (g) | 63.2 ± 11.9 | 57.3 ± 16.8 | 0.98 |
| Vitamin_C (mg) | 69.5 ± 25.5 | 74 ± 18 | 0.98 |
| Selenium (µg) | 55.5 ± 15.7 | 50.6 ± 16.5 | 0.99 |

